# The repressive genome compartment is established early in the cell cycle before forming the lamina associated domains

**DOI:** 10.1101/481598

**Authors:** TR Luperchio, MEG Sauria, VE Hoskins, X Wong, E DeBoy, M-C Gaillard, P Tsang, K Pekrun, RA Ach, NA Yamada, J Taylor, KL Reddy

## Abstract

Three-dimensional (3D) genome organization is thought to be important for regulation of gene expression. Chromosome conformation capture-based studies have uncovered ensemble organizational principles such as active (A) and inactive (B) compartmentalization. In addition, large inactive regions of the genome associate with the nuclear lamina, the Lamina Associated Domains (LADs). Here we investigate the dynamic relationship between A/B-compartment organization and the 3D organization of LADs. Using refined algorithms to identify active (A) and inactive (B) compartments from Hi-C data and to define LADs from DamID, we confirm that the LADs correspond to the B-compartment. Using specialized chromosome conformation paints, we show that LAD and A/B-compartment organization are dependent upon chromatin state and A-type lamins. By integrating single-cell Hi-C data with live cell imaging and chromosome conformation paints, we demonstrate that self-organization of the B-compartment within a chromosome is an early event post-mitosis and occurs prior to organization of these domains to the nuclear lamina.

## Introduction

DNA is highly and dynamically organized within the eukaryotic cell nucleus. This spatial organization has been implicated in a variety of crucial processes including sequestration of proteins involved in transcription, developmentally coordinated gene expression, and RNA processing and DNA repair into nuclear sub-domains. Nuclear organization manifests in a hierarchy of structures, each of which tends to favor self-interaction. At the whole-nucleus level, chromosomes occupy distinct regions in the nuclear volume called chromosome territories (CT), suggesting that each chromosome has a three-dimensional self-interacting organization^1–5^. These CTs are identifiable microscopically using whole chromosome-specific DNA probes or “paints” in a fluorescence in situ hybridization (FISH) assay. Subsequent FISH studies have demonstrated a sub-territory level of organization tightly linked to gene activity, with certain domains within a CT changing their relative disposition depending upon activity state^4^.

Recent high-throughput DNA sequencing based approaches, such as Hi-C or similar chromosome conformation capture (3C) based techniques, have been employed to uncover the organization of chromatin and identify self-interacting structures at the sequence level^6–9^. Two distinct types of structures have been found: locally self-interacting chromatin domains (TADs) and genome-wide predominantly bipartite spatial segregation known as the A-and B-compartment^7,8^. The B-compartment represents primarily repressed domains lacking self-interactions while the A-compartment displays robust self-interactions between active regions of the genome^8^. Compartment boundaries are typically bound by CTCF, which is highly depleted in the B-compartment, but can be found throughout the A-compartment in association with TADs^10^.

DNA Adenine Methyltransferase Identification (DamID) is a genome-wide technique to identify nuclear lamina-proximal chromatin, thus measuring the spatial distribution of chromosomal sub-domains within the nuclear volume^11–15^. These domains, termed Lamina Associated Domains (LADs) are approximately 100 kilobase (kb) to a megabase (Mb) in size and are enriched for transcriptionally silent genes and histone modifications indicative of facultative heterochromatin, such as histone H3 lysine 9 di-and trimethylation (H3K9me2/3) and histone H3 lysine 27 trimethylation (H3K27me3)^11,13,16–21^. Moreover, recent studies have demonstrated that both H3K9me2/3 and H3K27me3 are involved in LAD organization^13,18,19,22^. These findings, along with direct comparisons to Hi-C data and A/B compartment organization, have confirmed that the LADs represent a repressive genome compartment^14,23–26^. LADs are immediately flanked by active promoters, highlighting the stark delineation between repressed LAD domains and adjacent active regions. These borders also show enrichment for CTCF, as is seen in the boundaries between chromatin compartments observed by Hi-C^7,9,11,27^. LADs are enriched in the B-compartment^8,28,29^, although a detailed exploration of the relationship between the domain architecture of LADs and chromosomal sub-domain organization, such as the A/B-compartments, is surprisingly lacking.

How LADs organize at the single-cell level is unclear. One study used live cell imaging of a cancer cell line and followed LADs from one cell cycle to the next, finding that only 30% of regions identified by DamID are lamina-proximal in any single cell^18^. While some of the LADs repositioned to the lamina with fidelity, others appeared to remain in the nuclear interior in the subsequent cell cycle. These data suggest that the organization detected by ensemble techniques, such as DamID, might obscure significant cell-to-cell variability. Another study employing single-cell DamID in a haploid cancer cell line demonstrated that there is some variability between individual cells with a substantial set of core LADs consistently maintained at the lamina^17^. LADs that showed an increased variability of lamina association in this cell line were enriched in developmentally regulated genes. However, many developmentally regulated loci, such as the Igh locus in pro-B cells, display association with the nuclear lamina in greater than 90% of cells, as measured by 3D-FISH, suggesting a robust interaction of developmentally programmed genes with the nuclear lamina in relevant cell types^13,30–36^. One caveat to these and many other FISH studies is that they largely rely on mapping a single or small number of loci within the nucleus^37^. Such an approach inherently misses some important information—particularly the relationships of LADs with each other and their positioning relative to other LADs, non-LADs, and the nuclear lamina within the context of the entire chromosome polymer. While oligopaint technologies have been employed to identify the disposition of multiple locations within the nuclear volume, these approaches have relied on either assaying relatively small regions (up to several megabases), or entire chromosomes but with low coverage^38–40^.

Here we address the question of how LADs spatially organize within the nucleus, examining the relationships of LAD organization relative to both the nuclear lamina and non-LAD regions within an individual chromosome. We also investigate these relationships and spatial dynamics as the genome reorganizes post-mitosis and uncover an uncoupling of lamin association and LAD organization in early G1. In order to directly assess these relationships, we use two novel approaches to directly visualize LAD spatial organization at the single cell level: high density chromosome paints that differentially label LADs and non-LADs across an entire chromosome and a modified live cell LAD labeling system. Using high-density pools of chemically synthesized oligomers (Oligo Library Synthesis, OLS, Agilent Technologies) derived from DamID data in mouse embryonic fibroblasts (MEFs)^40,41^, we are able to detect LAD and non-LAD domains in situ and within the context of an entire chromosome. With this, we demonstrate a functional organization of the chromosome territory and epigenetic requirements for organization in single cells only hinted at by population-based assays such as Hi-C, ChIP and DamID. We also utilize our chromosome conformation paints to examine LAD dynamics through specific stages of the cell cycle. We show the distinct phases of genome partitioning that underlie A/B-compartmentalization, in which LADs first aggregate followed by movement to the periphery. We employ a LAD labeling approach in live cells, enabling the visualization of LAD self-aggregation post-mitosis and establishment of the chromosomal subdomain at the nuclear periphery in real time. Finally, we confirm these temporal dynamics through single-cell Hi-C. Taken together, we demonstrate that single-cell measures are indispensable for understanding spatial organization and dynamics of LADs.

## Results

### LADs, which largely correspond to the B-compartment, are reproducibly constrained in a peripheral zone of the nucleus in fibroblasts

To explore spatial organization and chromosome folding in single cells, we developed high resolution chromosome conformation oligopaints distinguishing LADs and non-LADs via a two fluorophore system. We first derived lamina-chromatin interaction maps in MEFs using DamID, which employs a bacterial adenine methyltransferase protein coupled to the nuclear lamina protein lamin B1 (LmnB1) (Fig. 1a)^11^. DamID material was initially hybridized to high-density tiling microarrays and replicate experiments were subsequently deep sequenced^13,42^. The resulting maps largely agreed with those previously published for MEFs (Supplementary Fig. 1). Using LAD regions defined by our DamID array maps in MEFs, we derived 150 base oligonucleotide probes for both LAD and non-LAD regions across chromosomes 11 and 12 separately, taking into account GC content, hybridization temperature and filtering for uniqueness of sequence in the genome. The resulting chromosome conformation paints were comprised of 0.5 million high-density oligonucleotide probes per chromosome and were divided into LAD and non-LAD pools that were chemically coupled to easily distinguishable fluorophores (Supplementary Fig. 1; Agilent Technologies).

**Figure 1:**
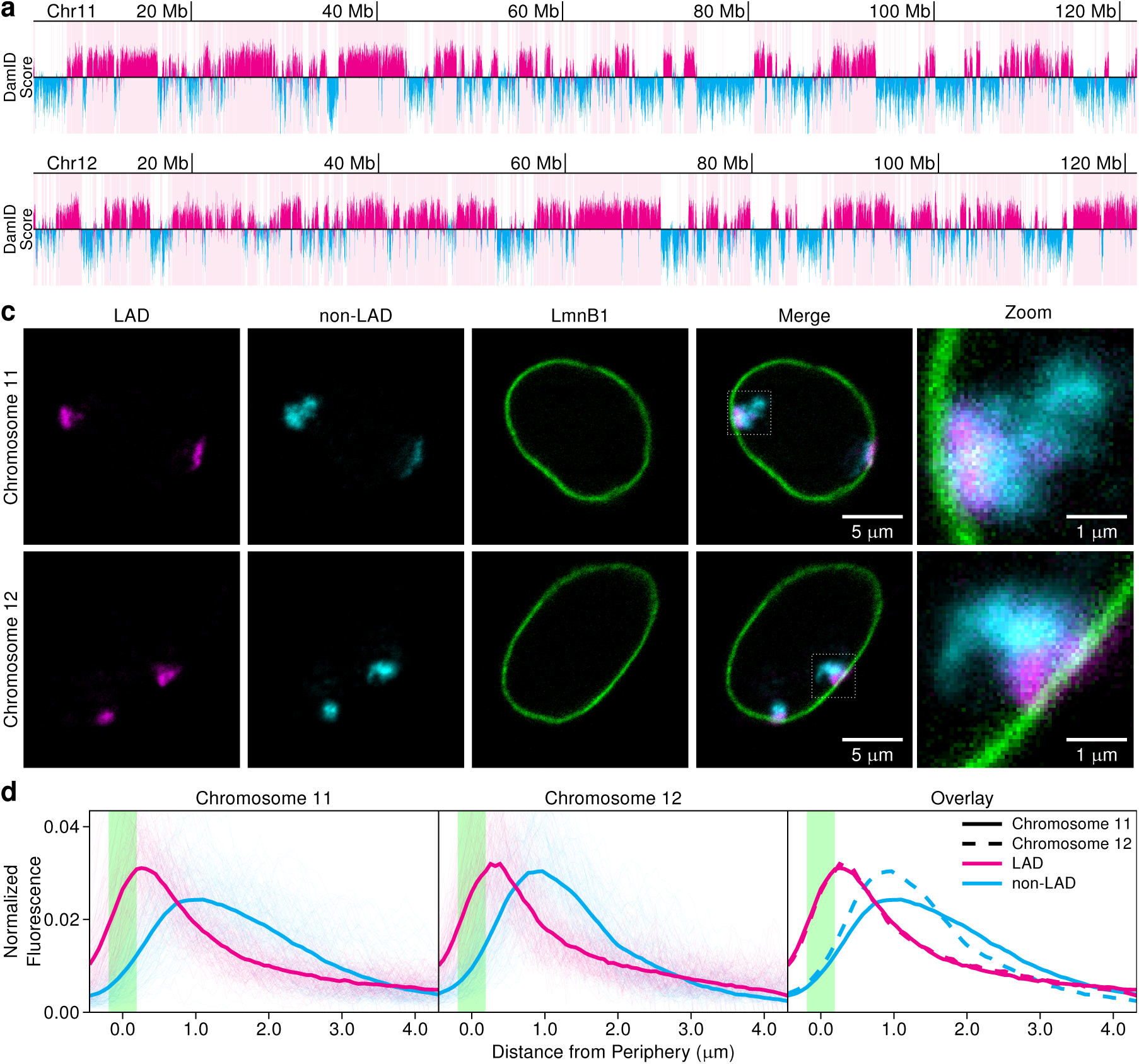
LAD definition and design of novel sub-chromosome compartment oligonucleotide paints. **a)** LmnB1 DamID log_2_ ratio plots for chromosomes 11 and 12 and LADs (solid pink bars) called by LADetector. **b)** 3D-immunoFISH probes (chromosome conformation paints) in single primary wild-type MEF nuclei reveal chromosome organization and the presence of LAD and non-LAD subdomains for both chromosome 11 and chromosome 12. **c)** Continuous measurements for chromosome 11 (n=51) and 12 (n=50) plotted to show the distributions of the LAD (magenta) and non-LAD (cyan) signals, as measured from the lamina (green), single cell measurements are shown as thin lines, average as thick lines.

In order to test whether the in situ organization of LADs is stochastic, as previously reported, or displays a more reproducible and constrained configuration, we performed FISH on 3D-preserved nuclei in ex vivo expanded early pass primary MEFs using our chromosome conformation paints^18,43^. LAD and non-LAD domains were clearly spatially segregated across the majority of their volumes and LADs were preferentially oriented near the lamina (Fig. 1b). To assess the distribution of domains, we measured fluorescence intensity in medial image planes for at least 50 chromosomes along lines perpendicular to the nuclear periphery (as demarcated by LmnB1 staining) and passing through LmnB1, LAD and non-LAD signals, with three line measurements per territory (Fig. 1c and Supplementary Fig. 2). For both chromosomes tested, LAD distributions demonstrated a close proximity to the lamina with a peak density of LAD signal at approximately 0.25 microns away and the majority of LAD signals resided within 0.6um of the nuclear lamina^18^. Distributions of LADs for both chromosomes 11 and 12 were almost identical despite differing LAD composition (49.97% versus 62.17% for chromosomes 11 and 12, respectively) (Fig. 1c and Supplementary Figs. 3 and 4) suggesting that constraint at the nuclear lamina results from the physical properties of the interface and not from overall LAD content. We also observed that LADs aggregated and formed a compact sub-territory, while non-LAD distributions were much broader, peaking nearly a micron away from the lamina. Unlike the LADs, the non-LAD distributions varied between chromosomes suggesting a different type of structure for these domains. The compact nature of these LAD domains at the lamina irrespective of differential LAD density leads us to describe this region where LADs are restricted and interface with the edge of the nucleus as the “peripheral zone”^32^. The existence of the peripheral zone is supported by studies that show an enrichment or specific exclusion of chromatin factors proximal to the nuclear lamina^18,33,43^. Taken together, our observation of the peripheral zone with the restriction of the majority of the LAD signal and an under-representation of non-LAD signals suggests a functional nuclear domain.

**Figure 2:**
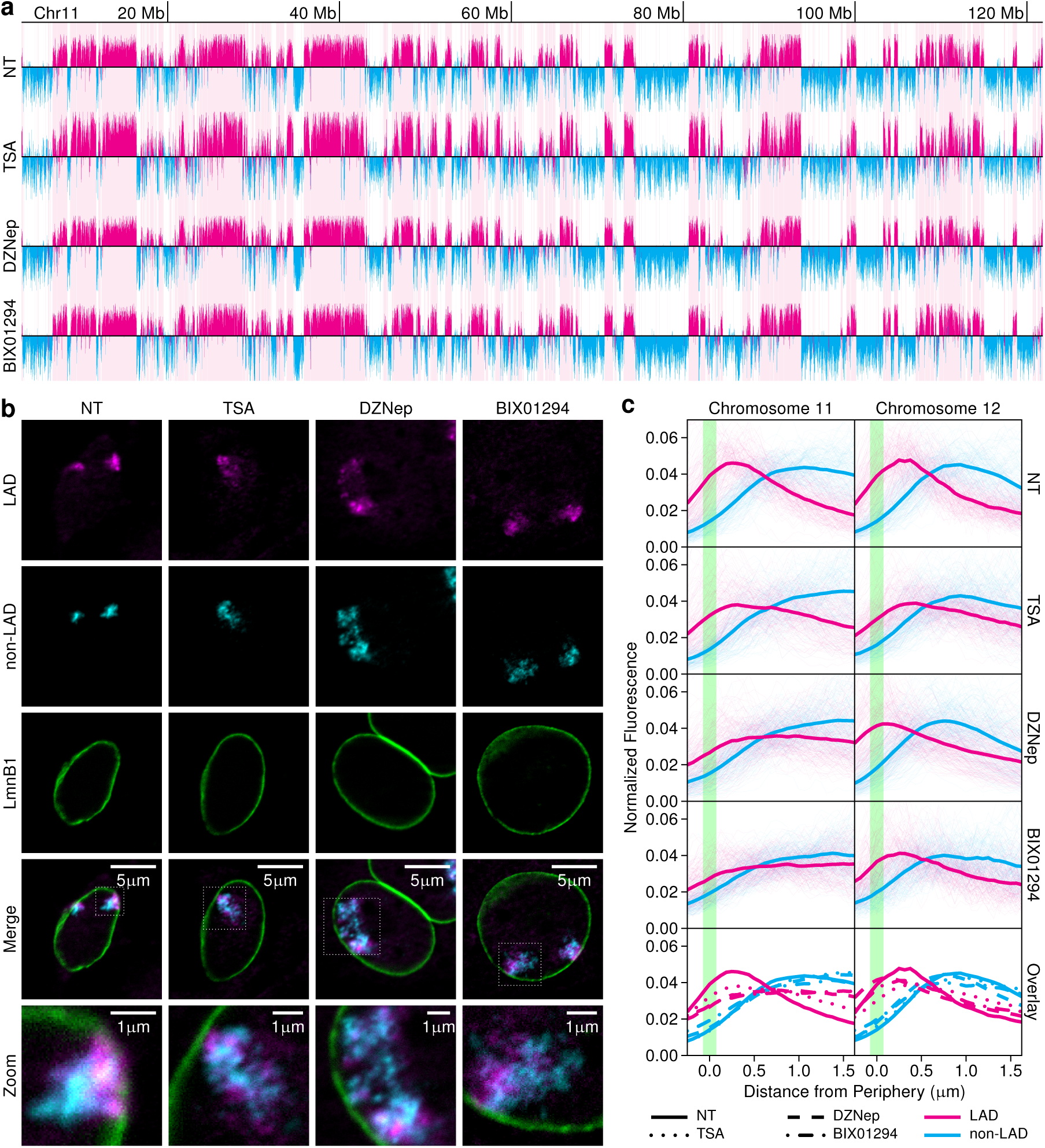
Epigenetic perturbation and sub-chromosomal architecture. **a)** Chromosome 11 LmnB1 DamID signal for primary non-treated (NT) and drug-treated MEF cells are shown with NT LAD calls indicated by pink bars and magenta signal. NT non-LAD signal is represented in cyan. **b)** 3D immunoFISH signals of LADs and non-LADs for non-treated cells and those treated with TSA, DZnep, or BIX01294 in chromosome 11 show perturbation of sub-territory organization. **c)** Individual measurements show distributions of LAD (magenta) and non-LADs (cyan) relative to LmnB1 (x=0, green). Individual measurements of chromosome territories are shown as thin lines for NT (Chr11 n=51, Chr12 n=50, see also Figure 1C), TSA treated (Chr11 n=52, Chr12 n=51), DZNep treated (Chr11 n=52, Chr12 n=54) and BIX01294 treated nuclei (Chr11 n=27, Chr12 n=15). Overlays of distributions for chromosomes 11 and 12 are provided in bottom two graphs.

**Figure 3:**
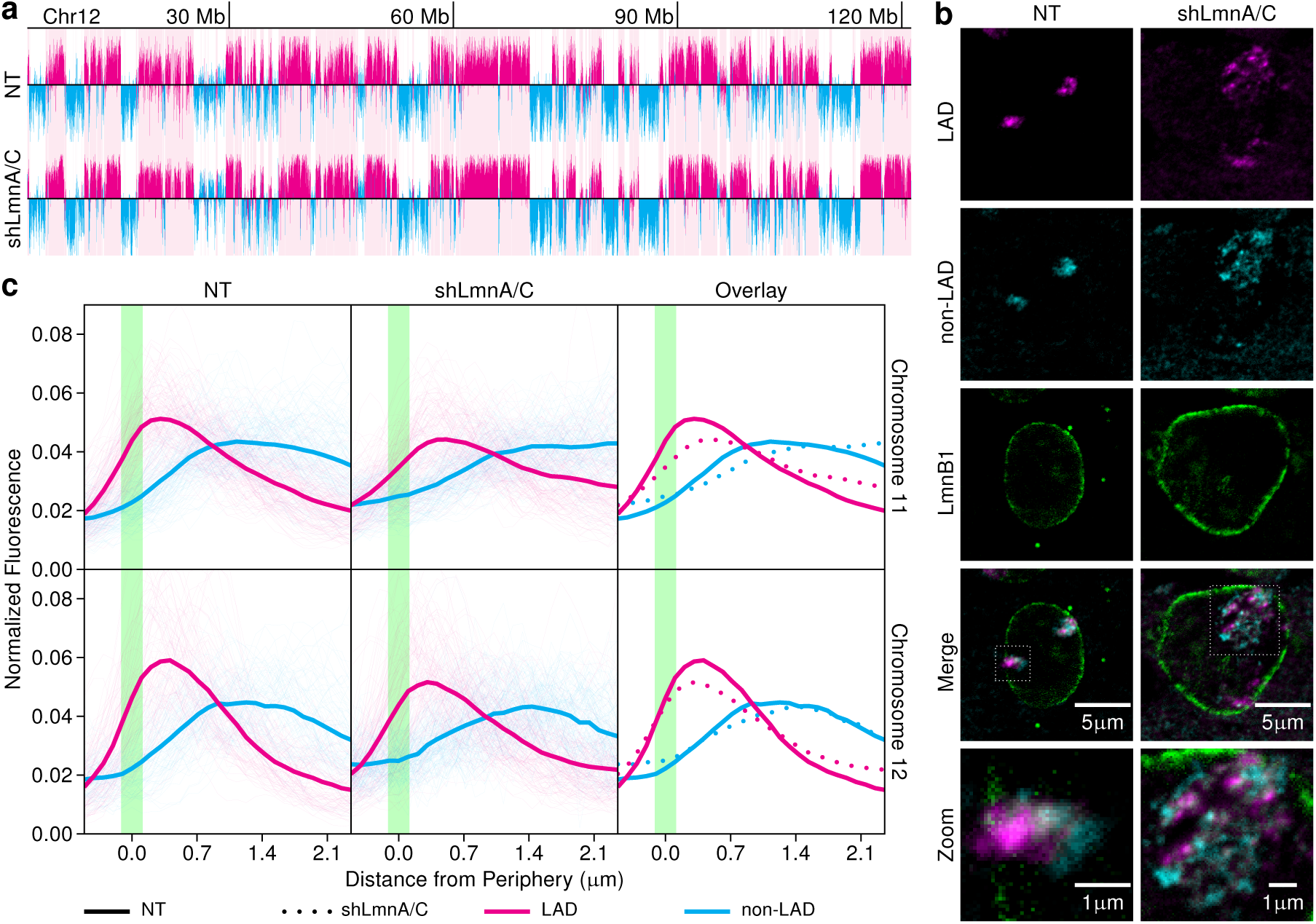
Nuclear structural integrity and sub-chromosomal architecture. **a)** LmnB1 DamID data for non-treated (NT) vs LmnA/C knockdown for chromosome 12. NT LAD calls are indicated by pink bars and magenta signal. NT non-LAD signal is represented in cyan. **b)** 3D immunoFISH signals of LADs (magenta) and non-LADs (cyan) in NT and LmnA/C knockdown highlight chromosome 12 sub-territory organization after LmnA/C knockdown. **c)** Continuous measurements for NT (Chr11 n=44, Chr12 n=28) and LmnA/C (Chr11 n=60, Chr12 n=67). Individual measurements show distributions of LAD (magenta) and non-LADs (cyan) relative to LmnB1 (x=0, green). Individual measurements of chromosome territories are shown as thin lines. Overlays of distributions for chromosomes 11 and 12 are provided in bottom two graphs.

**Figure 4:**
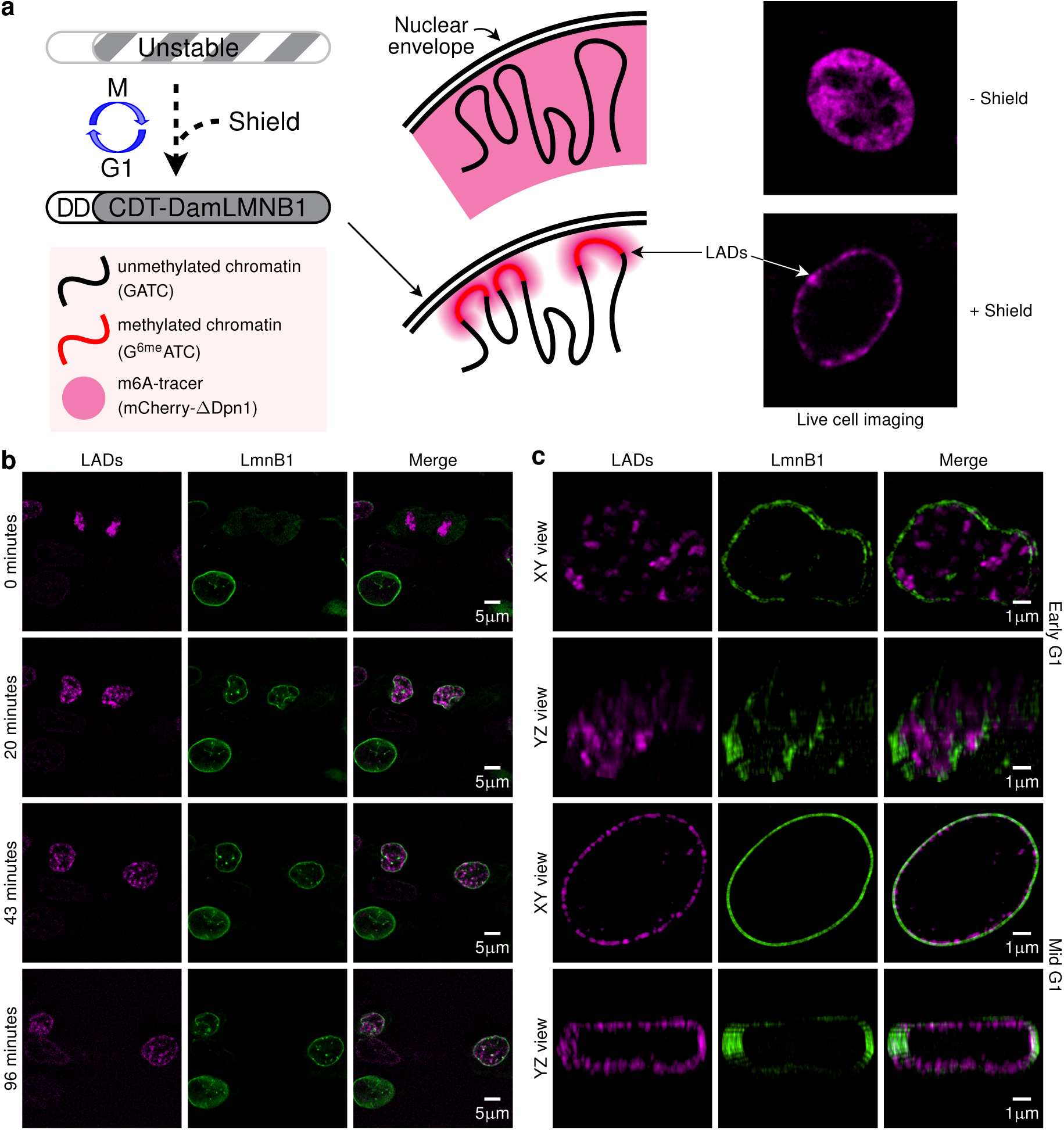
LAD self-aggregation occurs prior to peripheral localization, with lamina localization resolving by late G1. **a)** A modified m6A tracer system. Dam-LmnB1 construct containing the ubiquitination domain from cdt1 and a destabilization domain (DD) enables restriction of its stable expression to G1-phase of the cell cycle in the presence of a stabilizing reagent (Shield 1) for discrete labeling of adenines during G1. Similar to the previous system, a DpnI construct without a functional cleavage domain coupled to an mCherry (magenta) fluorophore allows visualization. Representative images + and - Shield. **b)** Live cell images of LADs/B compartment shown in magenta (m6A tracer), show a progression through early G1 for 96 min with the start of early G1 marked as 0 min. The nuclear periphery is shown in green using single chain antibody against LmnB1 (GFP-scfv Lamins) **c)** Super resolution microscopy of LADs using m6A tracer system in early G1 cell and mid G1 nuclei.

Previous studies have shown an enrichment of LADs in the B-compartment, suggesting that these designations may represent orthogonal measures of the same structures. In order to clarify the relationship between LADs and the B-compartment, we compared our genome-wide MEF LAD data with MEF Hi-C data using a refined compartment calling method^44,45^. To achieve a higher resolution compartment metric, we created a maximum likelihood-based score with independent distance-based signal decay curves depending on the compartment state of both interacting bins. LAD DamID showed a high degree of overlap with this Hi-C based compartment score, including a strong agreement between LAD and B-compartment state (85.5%) as well as boundary locations (Supplementary Fig. 5). Additionally, examination of B-compartment interactions showed self-associations within a single chromosome, which mirrors the compact organization of LADs of a single chromosome we observed by microscopy (Supplementary Fig. 5e,f). Taken together, these data suggest that LADs and the B-compartment are measures of the same structural domains. The regions of disagreement were highly enriched in ambiguous compartment scores (close to zero), representing regions of low Hi-C sequencing coverage, transient lamina association, or mixed state across the population. Because the chromosome conformation paints cover these LAD and non-LAD regions, they are able to highlight the spatial organization and segregation of the A- and B-compartments within a chromosome and act as a surrogate marker of the A- and B- compartments.

**Figure 5:**
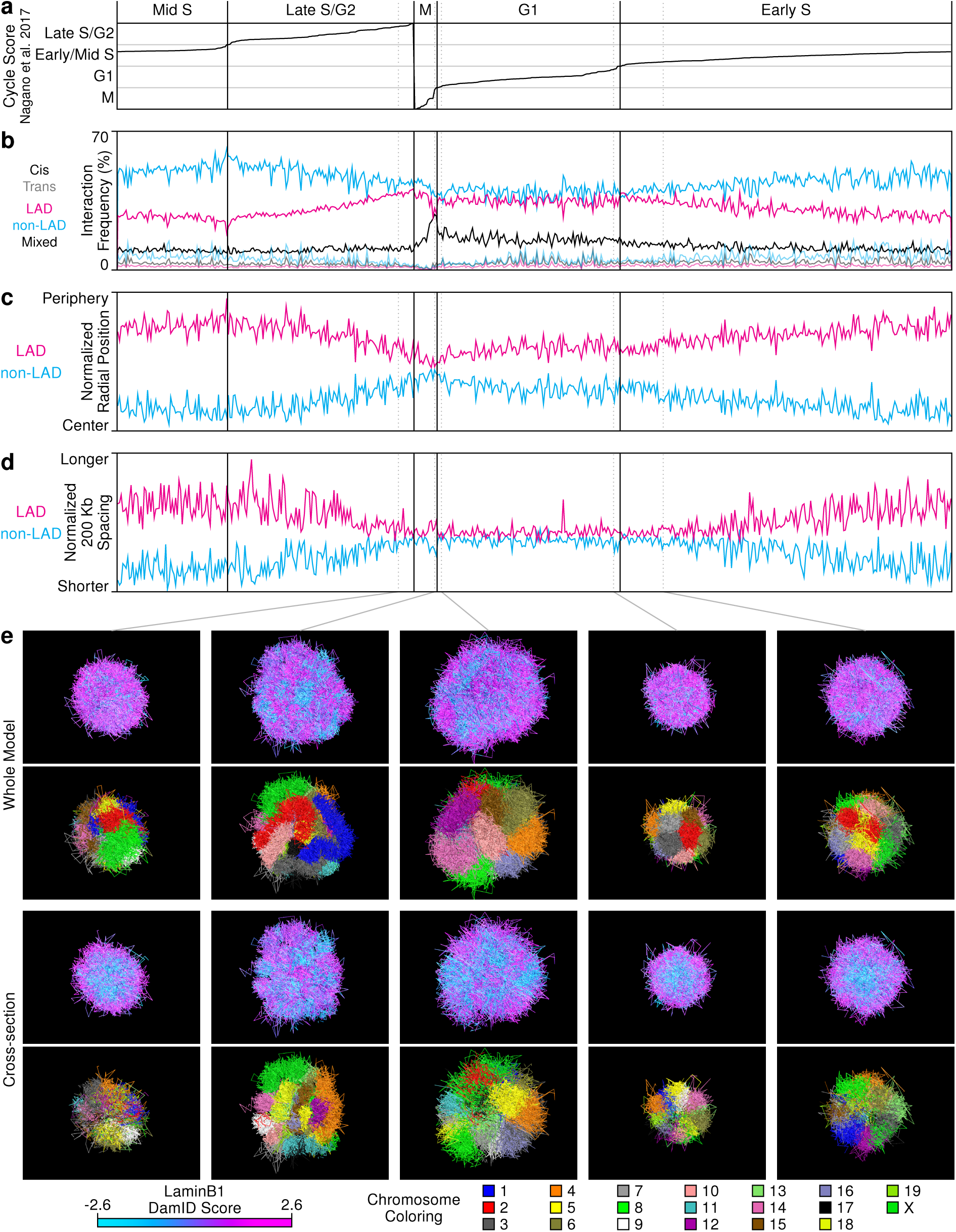
Modeling of single-cell Hi-C data. **a)** Cell cycle scores for high-coverage haploid cells taken from Nagano et al. (2017). **b)** Interaction frequency broken into groups by LAD/non-LAD and within chromosome (cis)/between chromosome (trans) features. **c)** Mean radial position of LAD and non-LAD 100Kb windows from scHi-C models, normalized by the mean radial position for all windows for each cell. **d)** Mean distance between sequence windows 200 Kb apart (midpoint to midpoint) and within the same LAD or non-LAD region as derived from the scHi-C models. Scores were normalized by the mean spacing across all 200 Kb-separated window pairs for each cell. **e)** Exemplar models for different phases of the cell cycle, colored to show either the DamID score (1st and 3rd rows) or indicate chromosome identity (2nd and 4th rows). The top two rows show whole models while the bottom two show cross-sections of the same models.

### LAD and A/B-compartment organization is dependent on both chromatin state and A-type lamins

Previous studies have demonstrated that the localization of at least some LADs to the nuclear lamina is dependent upon H3K9me2/3 and H3K27me3 and that the accumulation of these histone modifications may contribute mechanistically to LAD formation and maintenance^13,14,18,19,22^. Our group has previously shown that individual loci in LADs are directed away from the lamina upon disruption of either H3K9me2/3 or H3K27me3^13^. To test the impact of disruption of these epigenetic marks on chromosome organization as a whole, we treated primary MEFs with Trichostatin A (TSA, an HDAC inhibitor that promotes histone acetylation), BIX01294 (which inhibits H3K9me2 through inhibition of G9a and G9a-like protein) or 3-Deazaneplanocin A (DZNep, which decreases H3K27me3 through inhibiting EZH2)^13,22,46–48^. DamID showed little to no disruption of LAD organization by these measures with scores for DZNep, BIX01294, and TSA treatments having mean correlations across all replicate combinations of 90%, 89%, and 81% with non-treated cells, respectively, on par with the 89% correlation between non-treated replicates (Fig. 2a and Supplementary Fig. 6). These data appeared to contradict our previous study demonstrating the requirement for both H3K9me2/3 and H3K27me3 for individual LAD organization^13^. We hypothesized that since DamID is a longer-term ensemble measure of lamin association across a population of cells, single-cell variability in acute perturbations may be masked using this technique. To determine if these treatments altered in situ chromosome organization in single cells, we performed 3D-immunoFISH using our chromosome conformation paints (Fig. 2b). The level of disruption of LAD organization in individual cells was striking and varied from cell to cell (Supplementary Figs. 7 and 8). Many chromosomes could not be scored by our methodology because of the disruption and disaggregation of the sub-territories leading to a distribution of LADs through multiple planes of the nucleus or predominantly non-medial chromosome localization. These treatments also altered the morphology of the nuclei. In agreement with previous studies, disruption of H3K9me2/3 or H3K27me3 caused relocalization of some LADs away from the lamina, although some portion of LADs in both chromosome 11 and chromosome 12 remained proximal to the nuclear lamina. To remain consistent with the scoring of the untreated cells, scoring was performed only on chromosomes that displayed lamina proximal LAD signal in the medial planes and had 1-2 identifiable chromosome territories, leading to an under-representation of organizational disruption (Supplementary Figs. 7 and 8). Even with these limitations we were able to detect substantial reorganization (Fig. 2c). The most obvious effect of the drug treatments on LAD and non-LAD organization was dispersion and intermingling of both LAD and non-LAD chromatin, including a disaggregation of LADs that remained proximal to the lamina, suggesting a loss of A/B-compartmentalization and decrease in intra-chromosomal LAD associations. For all three drug treatments, our measurements demonstrate an expansion of the LAD sub-territory, loss of restriction to the peripheral zone, and an increase in the LAD and non-LAD spatial distribution variability across cells compared to untreated (Fig. 2c).

**Figure 6:**
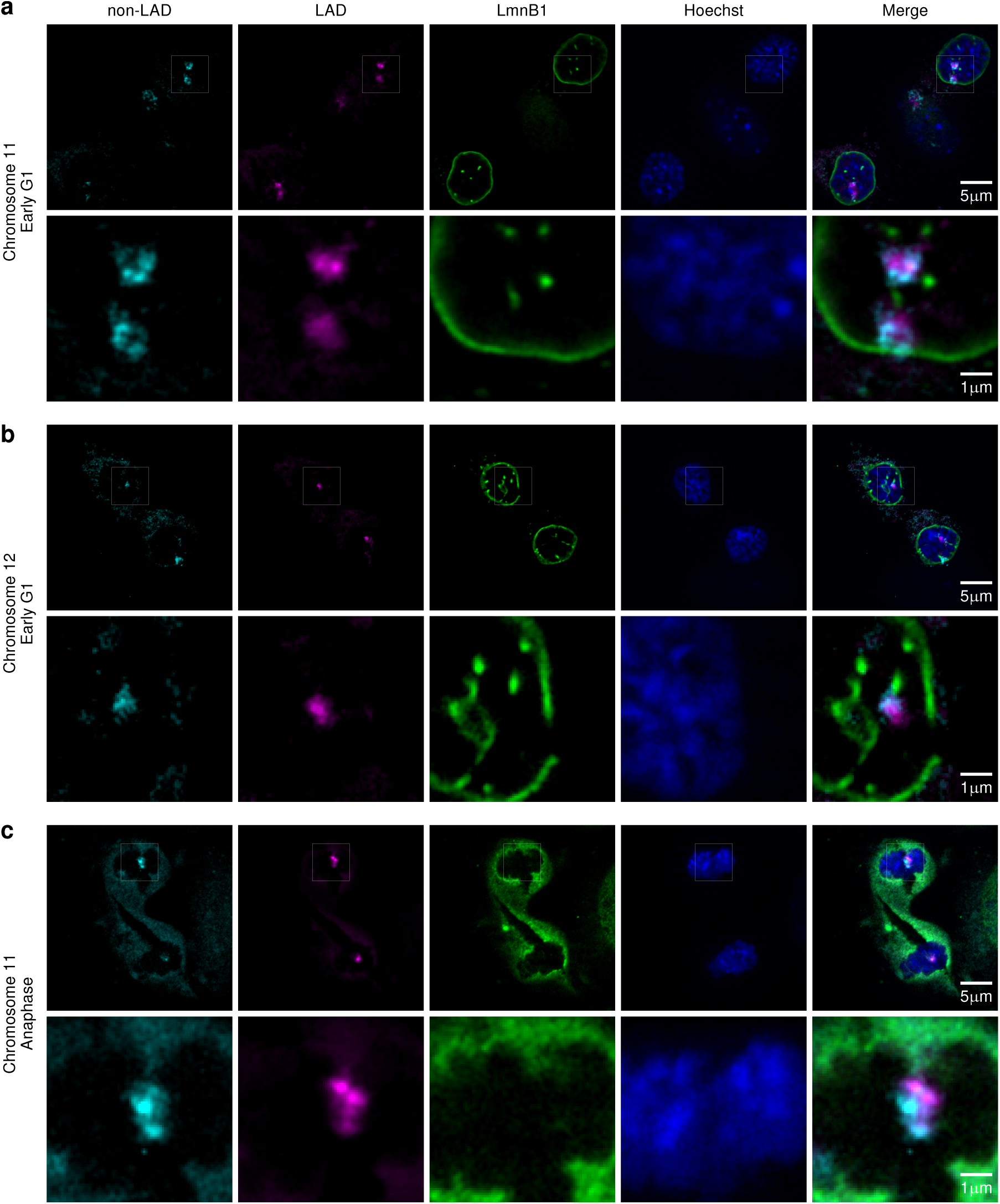
LAD/B-compartment aggregates are from single chromosomes. **a and b)** Chromosome conformation paints in primary wild-type MEF cells of early G1 for chromosomes 11 and 12 and **c)** chromosome 11 in an anaphase cell showing non-LADs (cyan), LADs (magenta), LmnB1 (green), and DNA (Hoechst).

**Figure 7:**
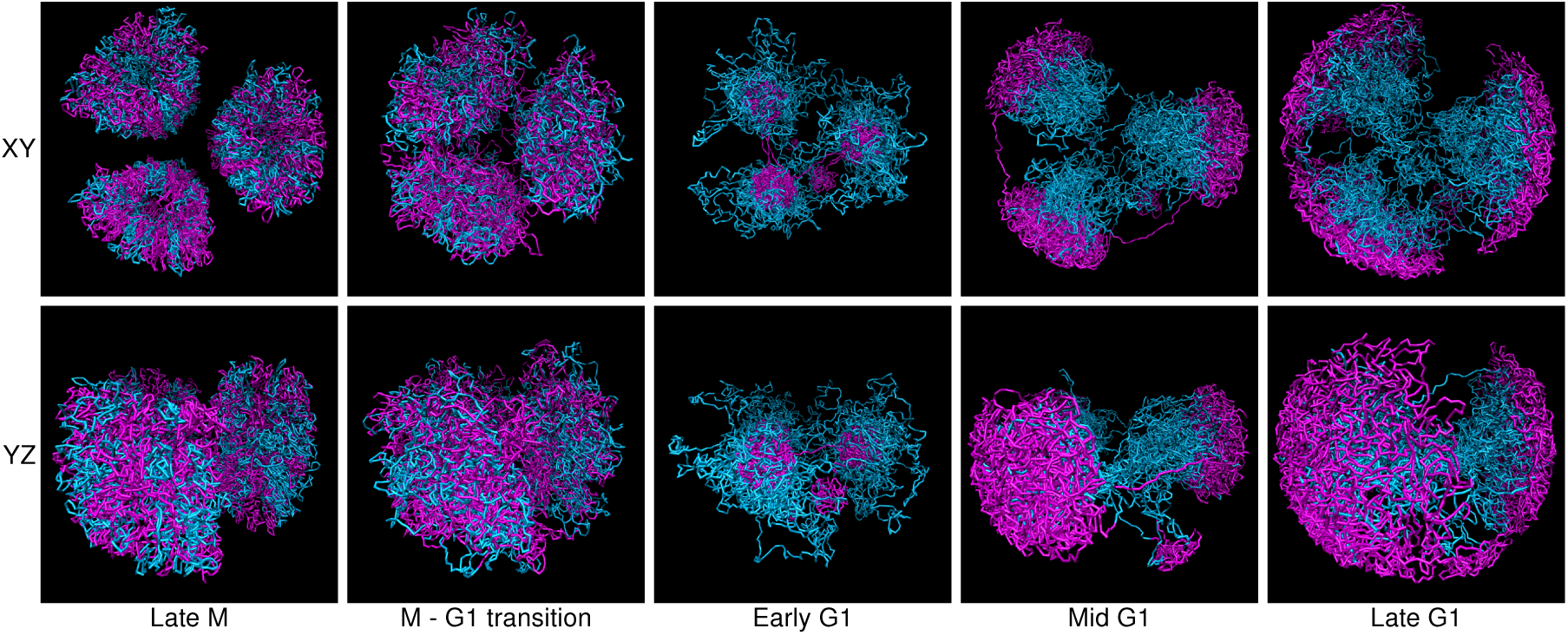
Model of compartment dynamics through mitosis and G1. A three chromosome cartoon nucleus demonstrating the proposed sequence of events following mitosis for the LAD (magenta)/non-LAD (cyan) spatial partitioning seen during the majority of the cell cycle. The two rows show a top and side view of the same process.

Previous data suggests a role for lamins in organizing chromatin to the nuclear lamina^13,49–51^. However, it is not clear how disruption of A-type lamins affects overall 3D organization within a chromosome. To test the role of lamin A/C (LmnA/C) in organizing LADs and the A/B-compartments across and within an individual chromosome, we removed LmnA/C by shRNA-mediated knockdown in MEFs. Knockdown of LmnA/C was nearly complete and did not affect either H3K9me2/3 or H3K27me3 (Supplementary Fig. 9a). Similar to what we observed for the epigenetic perturbations, loss of LmnA/C did not result in substantially altered LAD profiles by DamID (Fig. 3a). To examine if the DamID ensemble measure was missing perturbations at the single-cell level, we performed 3D-immunoFISH with the chromosome conformation paints. This revealed significant chromosome disorganization in individual cells (Fig. 3b,c). Because of the complete disruption to LAD organization, distribution throughout the nuclear volume, and loss of distinct chromosome territories, many nuclei could not be scored (Supplementary Fig. 9d,e). These disruptions included severe decompaction of the chromosome territory, intermingling of non-LAD and LAD signals (i.e. loss of A/B-compartmentalization), loss of peripheral association of many LADs and loss of LAD-to-LAD aggregation. We also observed that some of the organization of these chromosomes resulted in an “inversion” of chromosome organization, with LAD chromatin occupying the interior of the nucleus. This is in agreement with a previous study where an inversion of heterochromatin domains was noted in the absence of LmnA/C^49^. We note that this inverted chromatin phenotype was not observed for the epigenetic perturbations.

### The B-compartment/LADs display step-wise organization through the cell cycle

We have previously shown that reorganizing a de novo LAD region to the nuclear lamina requires transit through the cell cycle, suggesting a link between mitosis and dynamic LAD reorganization^14,15,49^. To explore the dynamics of LAD/B-compartment reconfiguration and LAD reestablishment to the nuclear periphery after mitosis, we modified the previously described m6A-tracer system that demarcates LADs in live cells to label LADs specifically in interphase (Fig. 4a)^18^.

By employing this modified LAD tracer system in a C57Bl/6j fibroblast cell line, along with a GFP-coupled single chain antibody to lamin, we observed that many LADs were not co-located with the reforming nuclear envelope during anaphase and telophase, in agreement with a previous study^18^. However, we found that these nucleoplasmic LADs did relocate to the nuclear periphery later in G1, taking nearly two hours post-mitosis to fully regain their lamina-proximal configuration (Fig. 4b). Based on the size and distribution of these LAD signals in early G1, we concluded that each aggregate was comprised of many LADs in the nuclear interior prior to lamina association. To measure the distribution and number of aggregates more carefully, we next employed super-resolution structured illumination microscopy (SIM) to measure LAD organization in mid-G1 versus early G1 cells (Fig. 4b,c). Each nucleus contained 70 to 100 LAD-aggregates. Accounting for ploidy (3-4N), this corresponded to 1-2 LAD aggregates per chromosome, suggesting that LADs within individual chromosomes self-aggregate. LAD signals at the periphery, while punctate in appearance, appeared to compress or spread out along the nuclear lamina later in G1.

### Separate compartmentalization and relocation observed in single-cell Hi-C data

To better understand the relationship between LAD organization and compartments during the cell cycle, we analyzed LAD organization within publicly available single-cell Hi-C (scHi-C) data obtained from cells throughout the cell cycle^52^. Our analysis used 506 haploid cell datasets spanning the entire cell cycle with a minimum of 100,000 interactions each. Cells were ordered through the cell cycle according to the scores detailed by Nagano et al. (Fig. 5a)^52^.

For every dataset, we ran multiple molecular dynamic annealing simulations at a resolution of 100 Kb to obtain three-dimensional chromatin configurations, selecting the best-fitting model for each cell. We then classified each 100 Kb region as LAD or non-LAD according to its majority membership. Our results confirm a dramatic reorganization of LADs and A/B-compartment configuration during different stages of the cell cycle and highlight the step-wise organization of LADs after mitosis. During mitosis, cells exhibited extensive interactions between LAD and non-LAD regions of chromatin, suggesting that A/B-compartmentalization is absent in this time-frame, in agreement with previous studies. Immediately after entering G1, we observed a large reduction in these inter-region interactions, while interactions within LAD chromatin, the presumptive B-compartment, increased (Fig. 5b), suggesting that compartmentalization is established almost immediately after mitosis, in strong agreement with our imaging data. This compartmentalization, also consistent with the imaging results, was not concurrent with the radial partitioning observed between non-LADs/LADs and A/B-compartment during the majority of the cell cycle (Fig. 5c). Instead, these data showed that LADs remained in the nuclear interior during early G1 as aggregated foci (B-compartment), reducing their contact with non-LAD chromatin (A-compartment). This radial segregation increased throughout G1 and S phase. We also examined local chromatin separation (distance between sequences 200 Kb apart) for both LADs and non-LADs (Fig. 5d). There was little difference in local separation of these domains at the resolution investigated here throughout G1. Differences only became evident in S phase, with relative LAD chromatin spacing increasing throughout until it peaked at the beginning of G2 and disappeared by mitosis. Taken together, the segregation of LADs at the periphery appear to involve three steps, which are supported by imaging data. First, a reduction in inter-state (A/B-compartment or non-LAD/LAD) interactions is accomplished independently of compaction or spatial isolation. Second, non-LADs/LADs are isolated from each other along the radial axis of the nucleus. Third, as LADs interact more with the periphery, individual LADs spread out across the lamina, suggesting that LAD/B-compartment regions assume a different topology compared to non-LAD/A-compartment regions. We also observed a reduction in the fidelity of chromosome territories as interphase proceeded (Fig. 5e). These appear to be driven entirely by non-LAD chromatin as the levels of LAD interchromosomal interactions remain unchanged throughout interphase.

### Chromosomes separate into subdomains independent of nuclear envelope assembly

Chromosome conformation paints and ensemble Hi-C data show that many LADs interact in a chromosome sub-territory, and both the LAD tracer system and single cell Hi-C data indicate that at least some LADs form an intra-chromosomal B-compartment prior to their organization to the lamina. By counting the number of large foci in the LAD-tracer system immediately post-mitosis, we inferred that the LADs may self interact in 1-2 large domains per chromosome. In order to determine if LADs from a single chromosome form a single aggregate and the relative organization of A-compartment chromatin, we used our chromosome conformation paints to visualize the LADs/non-LADs within either chromosome 11 or chromosome 12 transiting mitosis. In primary MEFs, early G1 daughter cells clearly showed a single LAD or B-compartment domain that was not associated with the nuclear lamina and was surrounded by the non-LAD or A-compartment chromatin, which was often found more proximal to the lamina at this stage (Fig. 6a,b). These data agree with formation of a single LAD/B-compartment aggregate within an individual chromosome. It is also clear that the nuclear lamina is not scaffolding the formation of the B-compartment or LADs, but rather that these interactions precede association with the nuclear periphery. These results highlight a transient stage in late anaphase, prior to nuclear envelope reformation but after daughter cell chromosome segregation, during which A/B-compartmentalization and LAD-LAD self-interactions are already being established (Fig. 6c).

## Discussion

Organization of LADs has been studied in a variety of cell types, both by genomic and cytological measures. Genome-wide approaches allow for global measurement of lamin association with high-resolution but give only an average measure of the population while single-cell approaches can capture variability between cells but at low resolution^17^. Conversely, cytological measures permit in situ measurements of nuclear partitioning either for specific loci in fixed cells (FISH), or LADs genome-wide in live cells (LAD-tracer system). With this study, we have bridged this gap using a combination of chromosome conformation paints and a modified LAD-tracer system to examine chromosome-wide LAD and A/B-compartment configurations in situ within the 3D context of the chromosome fiber and LAD dynamics through the cell cycle, respectively. Thus we establish organizing principles of LADs, both in their localization and formation.

Application of the chromosome conformation paints in primary early pass MEFs demonstrates that LADs are not strictly stochastic in their association with the nuclear lamina, but instead display a constrained configuration within a contiguously occupied peripheral zone (Fig. 1c). This zone results from the combination of lamina proximity and aggregation of LADs together, specifically those occupying the same chromosome. Our description of a peripheral zone compliments recent findings using TSA-seq as a molecular ruler defining an axis of increasing transcriptional activity from the lamina (repressive) to nuclear speckles (highly transcriptionally active)^24^. Our results also suggest that some LAD regions are more lamina distal, but still exist as part of a LAD aggregate seated against the lamina under spatial constraint. Using ensemble Hi-C measures, we confirm the within-chromosome aggregation of LADs and demonstrate the extensive overlap of LADs and the B-compartment, concluding our chromosome conformation paints allow us to visualize in vivo A/B-compartmentalization and both measures are indicative of LAD/B-compartment partitioning to the peripheral zone.

Our observations of LAD dynamics through the cell cycle further underscores LAD aggregation by showing it is independent of and precedes localization of the LADs to the nuclear lamina (Fig. 4). This process is also independent of the nuclear envelope, since aggregation of LADs occurs in anaphase, prior to nuclear envelope reformation. We propose that LAD formation and disposition is a three stage process beginning immediately post-mitosis. First, as chromatids decondense, LADs begin a process of aggregation, eventually forming foci (1-2 per chromosome) by early G1 (Figs. 6 and 7). Second, these foci are localized to the nuclear periphery and third, are subsequently spread out and adopt a more extended configuration as a greater portion of the LAD aggregates interact with the lamina (Fig. 7). However, the dissociation of the LADs from the lamina during mitosis begs the question, by what mechanism are LADs moved to the periphery. More comprehensive visualization techniques such as we have described may yet provide an answer.

We demonstrate that both aggregation and localization organizational forces are dependent on epigenetics (Figs. 2 and 3). Disruption of histone methylation machinery and increased histone acetylation led to similar outcomes, with disaggregation of the LAD sub-territory at the periphery and movement of some LADs away from the lamina. Our observations of LAD aggregation and a peripheral zone that partitions the A and B-compartments, rather than a strict coupling of the LAD regions to the lamina, are supported by recent work suggesting that heterochromatin is sequestered via phase separation^53–55^. What is less clear is the specific role of LmnA/C in this process. Previous work showed that there is a developmental transition from utilizing Lamin B Receptor (LBR) to LmnA/C for lamina association and constitutive loss of these proteins led to an inverted chromatin organization with heterochromatic domains occupying the nuclear interior^49^. In this study, our modeling of chromosome folding behavior, in agreement with our imaging data, show that LAD/B-compartment heterochromatin domains drive self-organization and that scaffolding at the lamina is an independent force contributing to the radial position of chromosomal domains, but not self-association of LADs. Here we show that acute loss of LmnA/C by shRNA-mediated depletion caused a subset of MEFs, which do not express appreciable levels of LBR, to display an inverted LAD organization, in agreement with those prior studies identifying inverted heterochromatic structures in the absence of Lamin A. We note that this phenotype was not fully penetrant, likely reflecting the varying levels of LmnA/C remaining in these cells or the time-scale during which the cells were lacking A-type lamins. Through our separate analysis of epigenetic marks and composition of the nuclear periphery, we have begun to dissect the roles each play in 3D nuclear organization, and suggest that aggregation of LADs and positioning at the nuclear periphery are temporally and spatially distinct processes^10,11,13^.

Understanding principles of LAD and genome organization is dependent on our ability to resolve both temporal dynamics and cell-to-cell variability. Furthermore, measuring these processes will necessarily need to leverage tools across different time and resolution scales. To this end, we use live cell imaging, high-density oligonucleotide chromosome conformation paints and single cell Hi-C data to reveal multiple forces working in collaboration to establish the canonical LAD and B-compartmentalization in interphase nuclei. Specifically, our chromosome conformation paints, designed based on genome-wide molecular data, clearly indicate preferential LAD and non-LAD organization throughout the majority of interphase. These data were confirmed both by live cell imaging and modeling of single cell Hi-C data. Strikingly, live cell imaging experiments measuring genome reorganization post-mitosis demonstrate that LADs/B-compartment regions self-associate prior to association with the nuclear lamina and single cell Hi-C data support the idea of such a step-wise mechanism. Using chromosome conformation paints during anaphase, we reveal that the subdomain organization of the chromosomes is the underlying driving force for compartmentalization, prior to even nuclear envelope reformation. Taking these results together, this multi-approach study highlights the complexity of the underlying forces working in conjunction to establish and maintain nuclear compartmentalization and chromatin architecture.

## Materials and Methods

### Contact for reagent and resource sharing

Further information and requests for resources and reagents should be directed to and will be fulfilled by the Lead Contact, Karen Reddy.

### Generation and maintenance of primary murine embryonic fibroblast (MEFs)

For primary MEFs, wild-type eight-week-old C57BL/6j mice were bred and embryos were harvested at E13.5. Individual embryos were homogenized using a razor blade, and cells were dissociated in 3 mL 0.05% trypsin for 20 min at 37°C, then 2 mL of 0.25% trypsin was added and and incubated again at 37°C for 5 min. Cells were pipetted vigorously to establish single cells, passed through a 70 µm cell strainer, pelleted and then plated in 10 cm dishes and labeled as P0. MEFs were cultured DMEM High Glucose with 10% FBS, penicillin/streptomycin, L-glutamine and non-essential amino acids. Cells were cultured for no longer than 5 passages before harvesting for experiments. For initial DamID experiments, longer term-culture C57BL/6 MEFs were purchased from ATCC (American Tissue Culture Collection, CRL-2752) and cultured according to their established protocols, in medium containing DMEM High, 10% FBS, Penicillin/Streptomycin and L-glutamine.

### Drug treatments

Primary MEFs were cultured as described and were treated with epigenetic modifying drugs for 24-60 hours, as previously described^13^. Drugs were added to the media at the following concentrations and refreshed at 24 hour intervals: 40 ng/mL TSA (Sigma, 1952), 0.5 µM BIX01294 (Ryan Scientific, RYS-AF-0051), 0.25 µM DZNep (Cayman Chemical, 13828, batch 0443536-5). For 3D-immunoFISH experiments, MEFs were treated with inhibitors while grown on slides. Wildtype MEFs served as non-treated controls and individual conditions were done as independent experiments. For drug treatment combined with DamID, primary MEFs were treated for 18-24 hours with the specified inhibitor, prior to infection with DamID virus.

### Lamin A/C knockdown

shRNA-mediated LmnA/C knockdown was carried out as described previously. Specifically, virus for knockdowns was generated in HEK 293T/17 cells (ATCC CRL-11268) by co-transfecting VSV-G, delta 8.9, and shLmnA/C (Sigma, clone NM_001002011.2-901s21c, 5’-GCGGCTTGTGGAGATCGATAA-3’) or shluciferase (5’-CGCTGAGTACTTCGAAATGTC-3’) with Fugene 6 transfection reagent (Promega E2691). 10 mM sodium butyrate was then added to the transfected cells 3 hours post transfection for an overnight incubation at 37°C, 5% CO2. The transfection media containing sodium butyrate was removed the following day and the cells were washed with 1X PBS. Opti-MEM was then added back to the cells which were then incubated at 37°C, 5% CO2. Viral supernatant was collected every 12 hours up to 3 collections and the supernatant of all 3 collections were pooled. Primary MEFs were cultured as described and incubated overnight with shLmnA/C or shluciferase fresh viral supernatants supplemented with 4 µg/mL polybrene and 10% FBS for 12-14 hours. Fresh MEF media was then added to the cells after the virus was removed and selected with 10 µg/ml blasticidin. For DamID profiling, cells were infected with DamID virus 4 days post shRNA transduction and cultured for additional 48 hours.

### DamID Infection

DamID was performed as described previously^11,13–15,21^. Cells were either transduced with murine retroviruses or with lentiviruses harboring the Dam constructs. Self-inactivating retroviral constructs pSMGV Dam-V5 (Dam-Only) and pSMGV Dam-V5-LaminB1 (Dam-LaminB1) were transfected using Fugene 6 transfection reagent (Promega, E2691) into the Platinum-E packaging line (Cell Biolabs, RV-101) to generate infectious particles. These viral supernatants in DMEM complete media were used to directly infect MEF lines. Lentiviral vectors pLGW-Dam and pLGW Dam-LmnB1 were co-transfected with VSV-G and delta 8.9 into HEK 293T/17 packaging cells using the Fugene 6 transfection reagent in DMEM High glucose complete media (DMEM High glucose supplemented with 10% FBS, Penicillin/Streptomycin, L-glutamine). 10 mM sodium butyrate was added to the transfected cells 3 hours post-transfection and left overnight. The following day this media was removed and the cells were washed briefly with 1X PBS before Opti-MEM media was added. Supernatants containing viral particles were collected every 12 hours between 36-72 hours after transfection, and these collections were pooled, filtered through 0.45 µM SFCA or PES, and then concentrated by ultracentrifugation. For infection with retrovirus or lentivirus, MEFs were incubated overnight with either Dam-only or Dam-LmnB1 viral supernatant and 4 µg polybrene. Cells were allowed to expand for 2-4 days then pelleted for harvest.

### DamID protocol

MEFs were collected by trypsinization and DNA was isolated using QIAamp DNA Mini kit (Qiagen, 51304), followed by ethanol precipitation and resuspension to 1 µg/ul in 10 mM Tris, pH 8.0. Digestion was performed overnight using 0.5-2.5 µg of this genomic DNA and restriction enzyme DpnI (NEB, R0176) and then heat-killed for 20 minutes at 80°C. Samples were cooled, then double stranded adapters of annealed oligonucleotides (IDT, HPLC purified) AdRt (5 ′ -CTAATACGACTCACTATAGGGCAGCGTGGTCGCGGCCGAGGA-3 ′) and AdRb (5′ -TCCTCGGCCG-3′) were ligated to the DpnI digested fragments in an overnight reaction at 16°C using T4 DNA ligase (Roche, 799009). After incubation the ligase was heat-inactivated at 65°C for 10 minutes, samples were cooled and then digested with DpnII for one hour at 37°C (NEB, R0543). These ligated pools were then amplified using AdR_PCR oligonucleotides as primer (5′ -GGTCGCGGCCGAGGATC-3′) (IDT) and Advantage cDNA polymerase mix (Clontech, 639105). Amplicons were electrophoresed in 1% agarose gel to check for amplification and the size distribution of the library and then column purified (Qiagen, 28104). Once purified, material was checked for LAD enrichment via qPCR (Applied Biosystems, 4368577 and StepOne Plus machine) using controls specific to an internal Immunoglobulin heavy chain (Igh) LAD region (J558 1, 5′ -AGTGCAGGGCTCACAGAAAA-3′, and J558 12, 5′ -CAGCTCCATCCCATGGTTAGA-3′) for validation prior to microarray hybridization and/or sequencing.

### DamID-seq library preparation

In order to ensure sequencing of all DamID fragments, post-DamID amplified material was randomized by performing an end repair reaction, followed by ligation and sonication. Specifically, 0.5-5 µg of column purified DamID material (from above) was end-repaired using the NEBNext End Repair Module (NEB E6050S) following manufacturer’s recommendations. After purification using the QIAquick PCR Purification Kit (Qiagen, 28104), 1µg of this material was then ligated in a volume of 20 µL with 1µl of T4 DNA ligase (Roche, 10799009001) at 16°C to generate a randomized library of large fragments. These large fragments were sonicated (in a volume of 200µL, 10mM Tris, pH 8.0) to generate fragments suitable for sequencing using a Bioruptor® UCD-200 at high power, 30 seconds ON, 30 seconds OFF for 1 hour in a 1.5 mL DNA LoBind microfuge tube (Eppendorf, 022431005). The DNA was then transferred to 1.5 ml TPX tubes (Diagenode, C30010010-1000) and sonicated for 4 rounds of 10 minutes (high power, 30 seconds ON and 30 seconds OFF). The DNA was transferred to new TPX tubes after each round to prevent etching of the TPX plastic. The sonication procedure yielded DNA sizes ranging from 100-200 bp. After sonication, the DNA was precipitated by adding 20 µl of 3M sodium acetate pH 5.5, 500 µl ethanol and supplemented with 3 µl of glycogen (molecular biology grade, 20 mg/ml) and kept at −80°C for at least 2 hours. The DNA mix was centrifuged at full speed for 10 min to pellet the sheared DNA with the carrier glycogen. The pellet was washed with 70% ethanol and then centrifuged again at full speed. The DNA pellet was then left to air dry. 20 µl of 10 mM Tris-HCl was used to resuspend the DNA pellet. 1 µl was quantified using the Quant-iT PicoGreen dsDNA kit (Invitrogen, P7589). Sequencing library preparation was performed using the NEBNext Ultra DNA library prep kit for Illumina (NEB, E7370S), following manufacturer’s instructions. Library quality and size was determined using a Bioanalyzer 2100 with DNA High Sensitivity reagents (Agilent, 5067-4626). Libraries were then quantified using the Kapa quantification Complete kit for Illumina (Kapa Biosystems, KK4824) on an Applied Biosystems 7500 Real Time qPCR system. Samples were normalized and pooled for multiplex sequencing.

### LAD and non-LAD chromosome-wide probe design and labeling

LADs from murine embryonic fibroblasts were defined through the LADetector algorithm, and complementary regions to Chromosomes 11 and 12 were defined as non-LADs. Data provided Geo GSE56990. Centromeres were excluded, and LAD and non-LADs were repeat masked. Probes were selected in silico based on TM and GC content, and those with high homology to off target loci were specifically removed. 150 base pair oligos were chemically synthesized using proprietary Agilent technology and probes were labeled in either Cy3 or Cy5 dyes using the Genomic DNA ULS Labeling Kit (Agilent, 5190-0419). 40 ng of LAD and non-LAD probes were combined with hybridization solution (10% dextran sulfate, 50% formamide, and 2X SSC) then denatured at 98°C for 5 minutes and pre-annealed at 37°C.

### 3D-ImmunoFISH and immunofluorescence

3D-immunoFISH was performed as described previously^13,15,49^. Briefly, primary fibroblast cells were plated on poly-L-lysine coated slides overnight. Cells on slides were fixed in 4% paraformaldehyde (PFA)/1X PBS for 15 minutes, then subjected to 3-5 minute washes in 1X PBS. After fixation and washing, cells were permeabilized in 0.5% TritonX-100/0.5% saponin for 15-20 minutes. The cells were washed 3 times 5 minutes each wash in 1X PBS, then acid treated in 0.1N hydrochloric acid for 12 minutes at room temperature. After acid treatment, slides were placed directly in 20% glycerol/1X PBS and then incubated at least one hour at room temperature or overnight at 4°C. After soaking in glycerol, cells were subjected to 4 freeze/thaw cycles by immersing glycerol coated slides in a liquid nitrogen bath. Cells were treated with RNAse (100 μg/ml) for 15 min in 2X SSC at room temperature in a humidified chamber. DNA in cells was denatured by incubating the slides in 70% formamide/2X SSC at 74°C for 3 min, then 50% formamide/2X SSC at 74°C for 1 min. After this denaturation, cells were covered with a coverslip containing chromosome conformation paints in hybridization solution and sealed. After overnight incubation at 37°C, slides were washed three times in 50% formamide/2X SSC at 47°C, three times with 63°C 0.2X SSC, one time with 2X SSC, and then two times with 1X PBS before blocking with 4% BSA in PBS for 30-60 min in a humidified chamber. Slides were then incubated with anti-LmnB1 primary antibody (1:200 dilution; Santa Cruz, SC-6217) in blocking medium overnight at 4°C. Slides were washed three times with 1X PBS/0.05% Triton X-100 and then incubated with secondary antibody in blocking medium DyLight 488 (1:200 dilution; Jackson ImmunoResearch, 211-482-171) for 1 hour at room temperature. Post incubation, slides were washed three times with 1X PBS/0.05% Triton X-100, and then DNA counterstained with 1 μg/ml Hoechst. Slides were then washed, mounted with SlowFade Gold (Life Technologies, S36936).

### Live cell imaging

B6 3T3 cells were infected to stably expressing ddDam-LaminB1-CDT, scfv-LMN-GFP and m6A tracer. ddDam-LaminB1-CDT is a destabilized version of the previously described DamID construct that has incorporated the CDT domain from the Fucci system to ensure its expression is restricted to interphase7–9. scfv-LMN-GFP is a lentiviral reconstruction of the Lamin-chromobody single chain antibody from Chromotek, that recognizes nuclear lamins. The m6A-tracer is comprised of a catalytically inactive version of Dpn1, that retains its ability to bind DNA, in frame with an mCherry red fluorescent protein 8,10. For cell cycle experiments, these cells were grown in the presence of shield ligand (AOBIOUS, AOB6677), which stabilizes the ddDam-LaminB1-CDT, along with 1mM thymidine (Sigma) block for 24 hours to enable synchronization of cells at G1/S9. This arrest was followed by release into complete DMEM medium (DMEM hi glucose, +10%FBS, 100 U/mL Penicillin and 100 μg/mL Streptomycin) containing 25µM 2-Deoxycytidine for 4 hours (no shield). Cells were then blocked at G2/M by incubation by replacing media with complete media (no shield) containing 10uM R0-3306 (AOBIOUS, AOB2010)for 16-20 hours11. Cells were released from this block by washing 3 times with warm Fluorobrite DMEM +10% FBS with 100 U/mL Penicillin and 100 μg/mL Streptomycin. 1-2 hours after release cells were imaged every 1-2 minutes using a 3i spinning disc confocal microscope or GE-OMX SIM super resolution microscope for early G1 imaging (https://microscopy.jhmi.edu/index.htm). Interphase cells were not synchronized and were imaged every 1-2 minutes.

### FISH image acquisition and processing

Slides were imaged using a Zeiss Axiovert fitted with an ApoTome and AxioCam MRm Camera. Imaging was performed at 100x or 63x with an Apochromat oil immersion objective with an NA of 1.5 using Immersol 518. (check all these details). AxioVision software was used to acquire images and .zvi files were exported and processed in FIJI^56^. Chromosome territories were evaluated for nuclear position and attachments to the lamina. As all chromosome 11 and 12 territories were visually determined maintain some level of proximity to the lamina, territories were measured through LAD signals closest to the lamina (lamin B1 signal) in medial planes. The distribution of LAD and non-LAD signals was measured using line scans in triplicate from outside to inside the nucleus and histogram measurements of pixel intensity were acquired for each channel using FIJI. Nuclei that were polyploid for chromosome 11 or 12, exhibited damage or were not fully visible in the field were excluded from the analysis. For each measurement, maximum lamin B1 signal was set to x=zero and all distances are relative to this zero point. Distance measurements were averaged together and normalized by total pixel intensity (Normalized value = Pixel intensity/Sum of total pixel intensity). Data collected from experiments performed on different days were pooled.

### DamID-seq data processing

DamID-seq reads were processed using LADetector (https://github.com/thereddylab/pyLAD), an updated and packaged version of the circular binary segmentation strategy previously described for identifying LADs from either array or sequencing data (<https://github.com/thereddylab/LADetector) ^13,14^. For arrays, DamID array signal intensity data were lifted over to mm9 using the Galaxy converter tool, and then data from replicate arrays were averaged together^57–59^ and quantile normalized and smoothed with the preprocessCore R package^60^. DamID array data were analyzed using a and earlier version of LADetector (https://github.com/thereddylab/LADetector). For both sequencing and array DamID data, LADs separated by less than 25 kb were considered to be part of a single LAD. All other parameters were left at default values. LADs were post-filtered to be greater than 100 kb, complementary genomic regions to LADs were defined as non-LADs. BedGraphs were generated for array data visualization using bedtools genomecov^61^ and output from the pyLAD LADetector for sequencing data.

## CTCF data

A normalized CTCF pileup was downloaded from GEO supplementary files (GSM918743)^62,63^.

### Hi-C normalization

Raw sequences for MEF Hi-C data from Krijger et al.^64^ were obtained from GEO^62^. Read ends were aligned to the mouse genome build 9 using BWA mem version 0.7.12-r1039^65^ and default settings. Reads were kept if they met one of the following criteria: Each read end mapped to a single position; one end failed to map but the other end mapped to two positions falling in two different restriction fragments; both ends mapped to no more than two positions from different restriction fragments and the downstream position of one end occurred on the same fragment as the upstream position of the other end. All replicates were combined. Reads were processed and normalized using HiFive version 1.3.2^44^. A maximum insert size of 650 bp was used to filter reads. Fends were filtered to have a minimum of one valid interaction.

The data were normalized using the binning algorithm correcting for GC content, fragment length, and mappability. GC content was calculated from the 200 bp upstream of restriction sites or the length of the fragment, whichever was shorter. Mappability was determined using the GEM mappability function, version 1.315^66^. Mappability of 36-mers was calculated every 10 bp with an approximation threshold of six, a maximum mismatch of 1 bp, and a minimum match of 28 bp. For each fend, the mean mappability score for the 200 bp upstream of the restriction site, or total fragment size if smaller, was used. For normalization, only intra-chromosomal reads with an interaction distance of at least 500 kb were used. GC content and fragment length were partitioned into 20 bins each and mappability was partitioned into 10 bins. All parameter partitions were done such that together they spanned the full range of values and contained equal numbers of fends in each bin. All bins were seeded from raw count means and GC and length parameters were optimized for up to 100 iterations or until the change in log-probability was less than one, whichever was achieved first.

### Hi-C compartment scoring

Eigenvector-based compartment scores were calculated as previously described^45^. Enrichments were calculated for either 1 Mb (low-resolution) or 10 kb bins (high-resolution). Bins were expanded using HiFive’s dynamic binning to a minimum or 3 reads per bin. For each pairwise combination of rows for the enrichment heatmap, the Pearson correlation was calculated. Taking the first eigenvector of the correlation matrix yielded the eigenvector-based compartment score. Because the sign of the eigenvector is random, we used mean transcriptional activity in positively versus negatively scored regions to determine A and B-compartment score signs. Where necessary, signs were flipped so that all B-compartments corresponded to positive eigenvector scores.

Likelihood compartment scores were calculated as the log2-transformed ratio of the probability of each 10 kb interval occurring in the B-compartment divided by the probability of that interval occurring in the A-compartment. The sign of the high-resolution first eigenvector score described above determined compartment initialization (positive values were associated with the B-compartment). Bins with fewer than five interactions longer than 500 Kb were removed. Interactions spanning 500 Kb or greater were divided into three groups: both sides occurring in the A-compartment, the B-compartment, or one side in each compartment. The distance dependent signal curve for each category was calculated by find the sum of counts divided by the sum of expected values at each distance interval. For distance intervals containing fewer than 10,000 reads were joined with the next largest interval prior to finding enrichment. This was continued until the 10,000 read minimum was met. The effective distance for joined bins was calculated as the mean of the log-transformed bin distances. Enrichment values for distances corresponding to bins that had fewer than 10,000 reads were interpolated linearly based on the log-transformed expected values and log-transformed distances of the two adjacent bins. The probability for each interval was calculated under the Poisson distribution as follows:

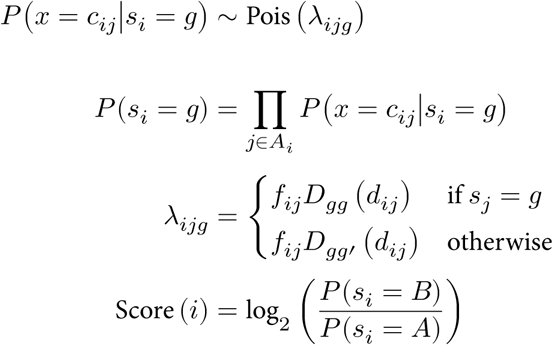

where *s*_*i*_ is the compartment state of interval *i, A*_*i*_ is the set of valid interactions of at least 500 Kb involving interval *i, c*_*ij*_ is the sum of observed counts for the interaction bin between intervals *i* and *j, f*_*ij*_ is the sum of interaction normalization values for bin *ij, d*_*ij*_ is the distance between midpoints of intervals *i* and *j*, and *D*_*gg*_ and *D*_*gg’*_ are the distance dependent signal functions for within compartments of type *g* and between different compartments, respectively.

Training was accomplished on a chromosome by chromosome basis in an iterative fashion, calculating the distance dependent signal curves, calculating the compartment scores, updating the top 50% of scores (rounding up), and adjusting states based on the signs of the scores. This was performed for up to 200 rounds. If a stable set of interactions was achieved, the associated scores were kept. If a chromosome began switching between two sets of stable states, the mean of these two sets of scores was taken. Otherwise after a 20 round burn-in period, scores were sampled every round and the mean score for each interval was taken after the final iteration.

### Single cell Hi-C modeling

Haploid single cell Hi-C processed counts from Nagano et al.^52^ were obtained from the Tanay lab (http://compgenomics.weizmann.ac.il/files/archives/schic_hap_2i_adj_files.tar.gz and http://compgenomics.weizmann.ac.il/files/archives/schic_hap_serum_adj_files.tar.gz). Only cells with a total of 100,000 reads or more were used. Data were further filtered using HiFive single cell Hi-C filters. This involved removing fragment ends (fends) with no interactions, fends smaller than 21 bp or larger than 10 Kb, and all fends not originating from chromosomes 1 through 19 or X. Next, because only haploid cell data were used any fend with more than two interactions was removed and fends with exactly two interactions were removed if the interactions occurred with partner fends more than 40 fends apart; otherwise, the longer of the two interactions was kept. Finally, fends were partitioned into 1 Mb bins and a connectivity graph was created with edges present if at least one unfiltered interaction existed between bins. For each edge, interactions were removed if the next shortest path between bins was longer than three steps. Modeling was performed for each cell dataset using a coarse grained annealing molecular dynamic simulation. Chromatin was represented as beads representing 100 Kb beads, starting from the first 100 Kb bin for each chromosome containing at least one valid interaction and ending with the last bin containing a valid interaction. Intervening bins containing no interactions were kept for the purposes of modeling but excluded for all subsequent analyses. The force field was setup similar to that described by Nagano et al.^67^. Two forces were applied to each bead, a general repulsive force and a harmonic bonding force. All pairwise combinations of beads, with the exception of those having scHi-C interactions, were given a repulsive force with a scaling factor (*k*_*1*_) of one for distances less than 60 nm (*d*_*lim*_).

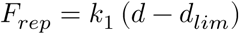

All beads representing adjacent chromatin bins (backbone) and bead pairs with scHi-C interactions (constraint) were given a flat-bottomed harmonic potential force with a scaling factor of 25 (*k*_*2*_) around an effective target distance (*d*_*eff*_) of 150 nm or 120 nm (*d*_*target*_) for backbone and constraint bonds, respectively, scaled by the inverse of the square-root of the number of observed valid reads (*r*) supporting an interaction (backbone bonds were always given a distance scaling factor of one). At distances less than 15% of *d*_*target*_ (*d*_*lower*_) for constraint bonds (there was no lower limit for backbone bonds, thus *d*_*lower*_ equaled 0), an exponential repulsive force was applied. At distances between *d*_*target*_ and *d*_*target*_ + 30 nm (*d*_*upper*_), an exponential attractive force was applied. At distances greater than *d*_*upper*_, the attractive force became linear.

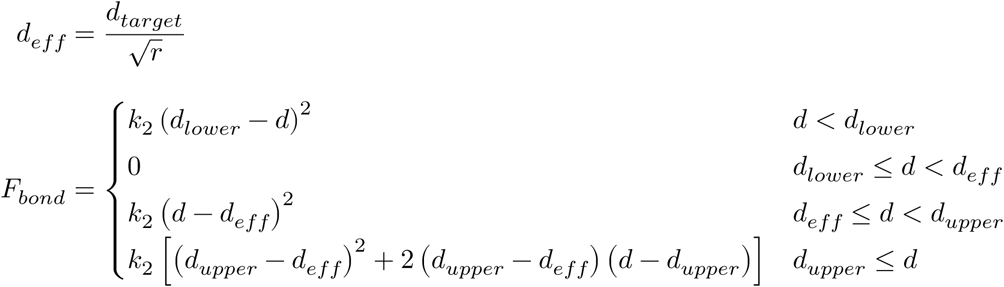

Simulations were run using OpenMM version 7.1.1 and wrapped using the Mirny lab’s openmm-polymer (https://bitbucket.org/mirnylab/openmm-polymer). Specific forces were custom implemented and available in the accompanying code. Initial model conformations generated by randomly ordering chromosomes, end to end, and treating all beads as a single polymer. Beads were arranged around a 3 µm circle, evenly spaced along 5 oscillations of a sine wave perpendicular to the plane of the circle with an amplitude of 1.5 µm. Each simulation was run for 301 time steps, each consisting of 1000 motion steps. For the first 101 time steps, the temperature was linearly ramped from 10000 K to 5000 K. At the same time, the *k*_*2*_ parameter for constraint bonds was ramped from 0 to 25. During the remaining 200 time steps, the temperature was linearly ramped down from 5000 K to 10 K. Each simulation was repeated 10 times and the resulting model with the fewest bonds exceeding their effective target lengths was selected for downstream analysis.

For each scHi-C cell model, a hull was constructed based on polymer bead positions. Initially, for each chromosome, the set of distances for that chromosome’s beads from the chromosome’s mean bead position was calculated. The standard deviation across all sets of chromosome distances was determined and any beads whose distance exceeded 3 standards of deviation was removed from the set used to determine the nuclear hull. For each chromosome, the valid beads were used to construct a convex hull. The nuclear hull was defined as the union of all chromosome hulls.

For each bead, a ray was projected from the hull center of mass, through the bead position. Next, the longest distance from the center of mass position to an intersection with the nuclear hull was found. Because nuclear hulls were not guaranteed to be convex, the ray could intersect the hull multiple times. Only the furthest distance was used. The radial position was defined as the bead distance from the center of mass divided by the projection intersection distance from the center of mass. In the case of outlier beads excluded from the hull-defining set, if the bead distance exceeded the projection distance, the radial position was defined as one.

Beads were assigned a LAD/non-LAD state based on LAD calls from Peric-Hupkes et al. (GEO ID GSM426758)^68^, where the bead was assigned the state that corresponded to the majority of its bases. Model compaction was determined from scHi-C models by examining every chromosome bead triplet and if all three beads were in the same state (in or out of a LAD), calculating the distance between the end beads.

## Author Contributions

T.R.L., V.E.H, X.W., and K.L.R. designed the project, T.R.L., X.W., T.H., E.D. M-C.G., M.E.G.S and K.L.R. executed the experiments, P.T., R.A.A., K.P., and N.A.Y. designed the oligonucleotide pools for paints, M.E.G.S., T.R.L., X.W. and J.T. analyzed and interpreted the data, K.L.R. and M.E.G.S. wrote the manuscript with input from J.T. and T.R.L. All authors edited and approved the final manuscript.

## ACKNOWLEDGEMENTS

We thank the Reeves lab for help making MEFs. We thank Rakel Tryggvadottir and Sinan Ramazanoglu for invaluable help with sequencing. We thank the JHU Provost’s Office for support through the Catalyst and Discovery Awards. T.R.L. was funded from NIH Training Grant T32GM007445, K.L.R and M-C.G. were partly funded from R21AG050132, and M.E.G.S and J.T. were funded in part by NIH/NIDDK grant R24 DK106766 and NIH/NHGRI grant U41 HG006620.

## Supplemental Figures

**Supplemental Figure 1:**
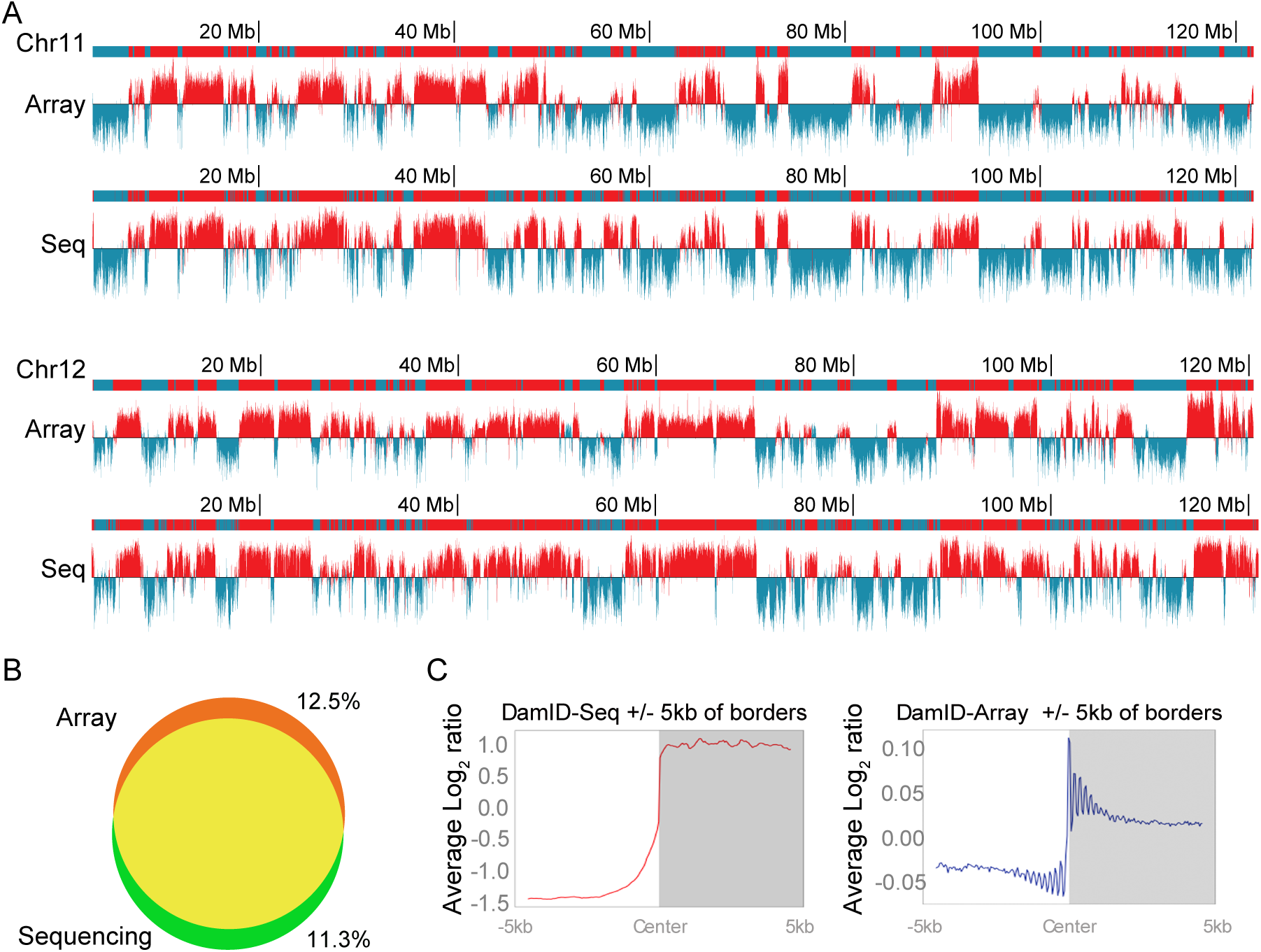
Comparison of DamID-array and DamID-seq. **a)** Lamina-chromatin contact maps derived from DamID-array (top) or DamID-seq (bottom) for chromosome 11 and 12 arranged for visual comparison. **b)** Venn diagram features overlap of LADs defined by DamID-seq and DamID-array, <13% are unique between techniques, similar to differences observed between replicate experiments (see text). **c)** Plots of the average log2 ratios of DamID-seq signals (red line) and DamID-array signals (blue line) outside (white box) and inside (gray box) LAD regions. Region shown is +/-5 kb of a LAD border.

**Supplemental Figure 2:**
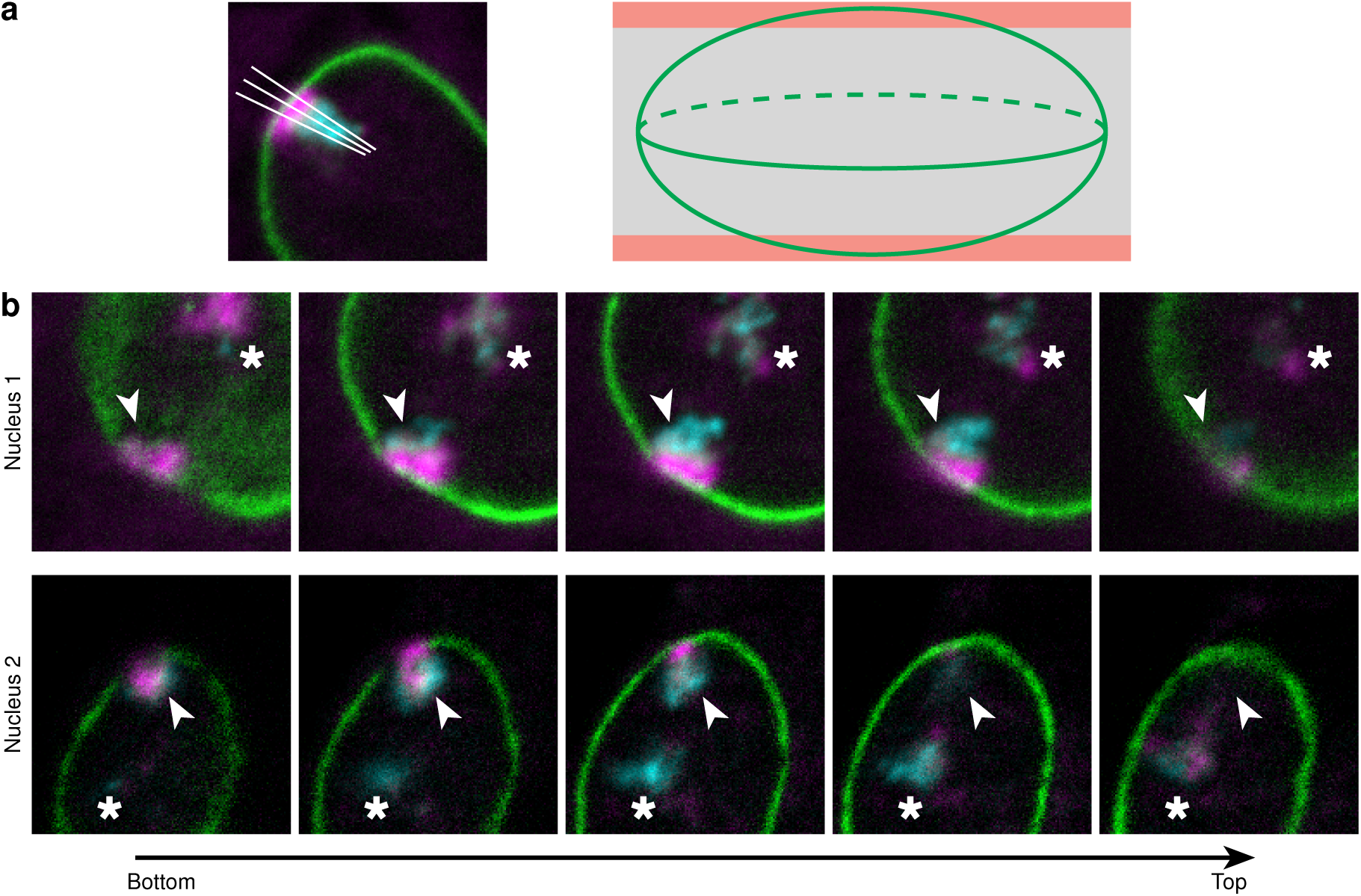
Scoring methodology. **a)** Line scan measurements to collect distribution of LAD and non-LAD sub-territories from the lamina were done by collecting three measures across the chromosome territory, from the lamina through the chromosome territory, passing through the shortest distance of lamin signal to LAD signal. Measurements were taken in the medial planes, highlighted gray shaded area of nucleus schematic. **b)** Examples of territory disposition in the nuclear volume are shown. Two nuclei are presented as 5 slices, from top toward the bottom of the nucleus. The territory that was scored in medial planes is indicated with an arrow head. Territory that was not scored is starred, because the majority of the LAD signal was at the top or bottom of the nucleus. Line scans were done in single or multiple planes, depending upon the disposition and intensity of LAD signals.

**Supplemental Figure 3:**
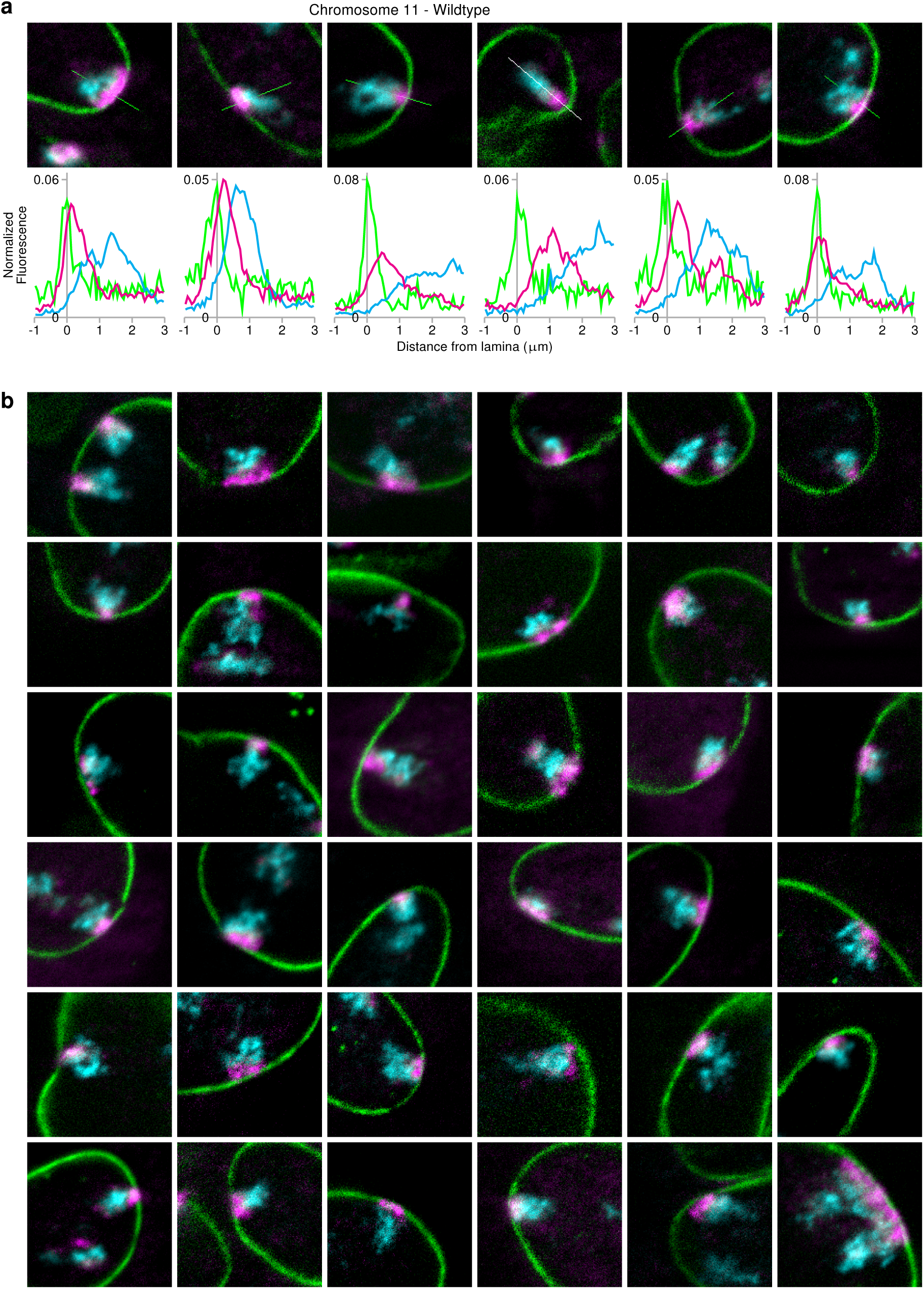
Chromosome conformation paints for chromosome 11. **a)** Examples of single line scan measurements (line overlaid on chromosome territory) with accompanying plot profiles (graphs, below) for LADs (magenta) non-LADs (cyan) and LmnB1 (green). **b)** Array of chromosome 11 territories visualized by chromosome conformation paints. All images are shown at the same magnification. All graphs include measurements to 3 µm from the lamina.

**Supplemental Figure 4:**
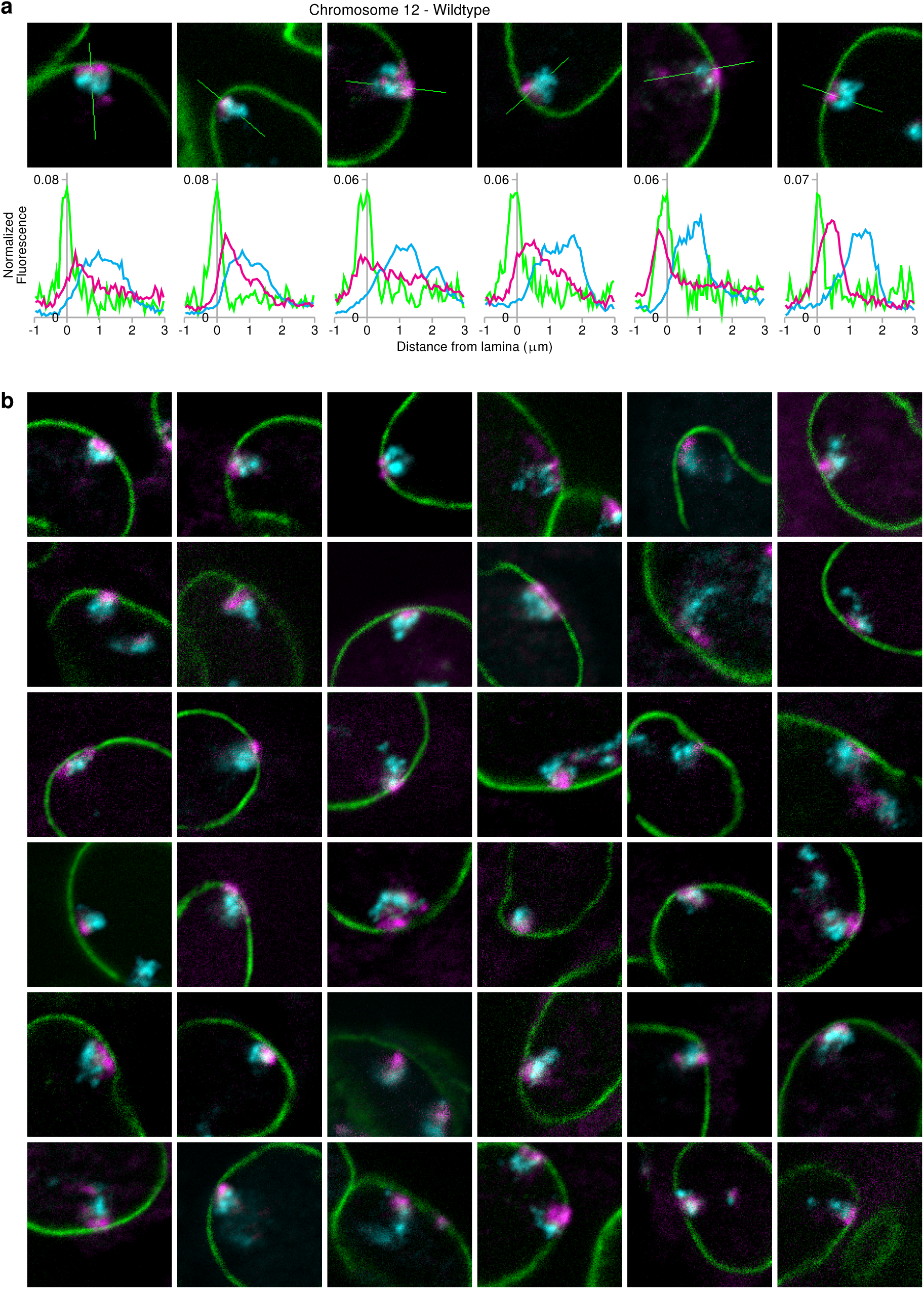
Chromosome conformation paints for chromosome 12. **a)** Examples of single line scan measurements (line overlaid on chromosome territory) with accompanying plot profiles (graphs, below) for LADs (magenta) non-LADs (cyan) and LmnB1 (green). **b)** Array of chromosome 12 territories visualized by chromosome conformation paints. All images are shown at the same magnification. All graphs include measurements to 3 µm from the lamina.

**Supplemental Figure 5:**
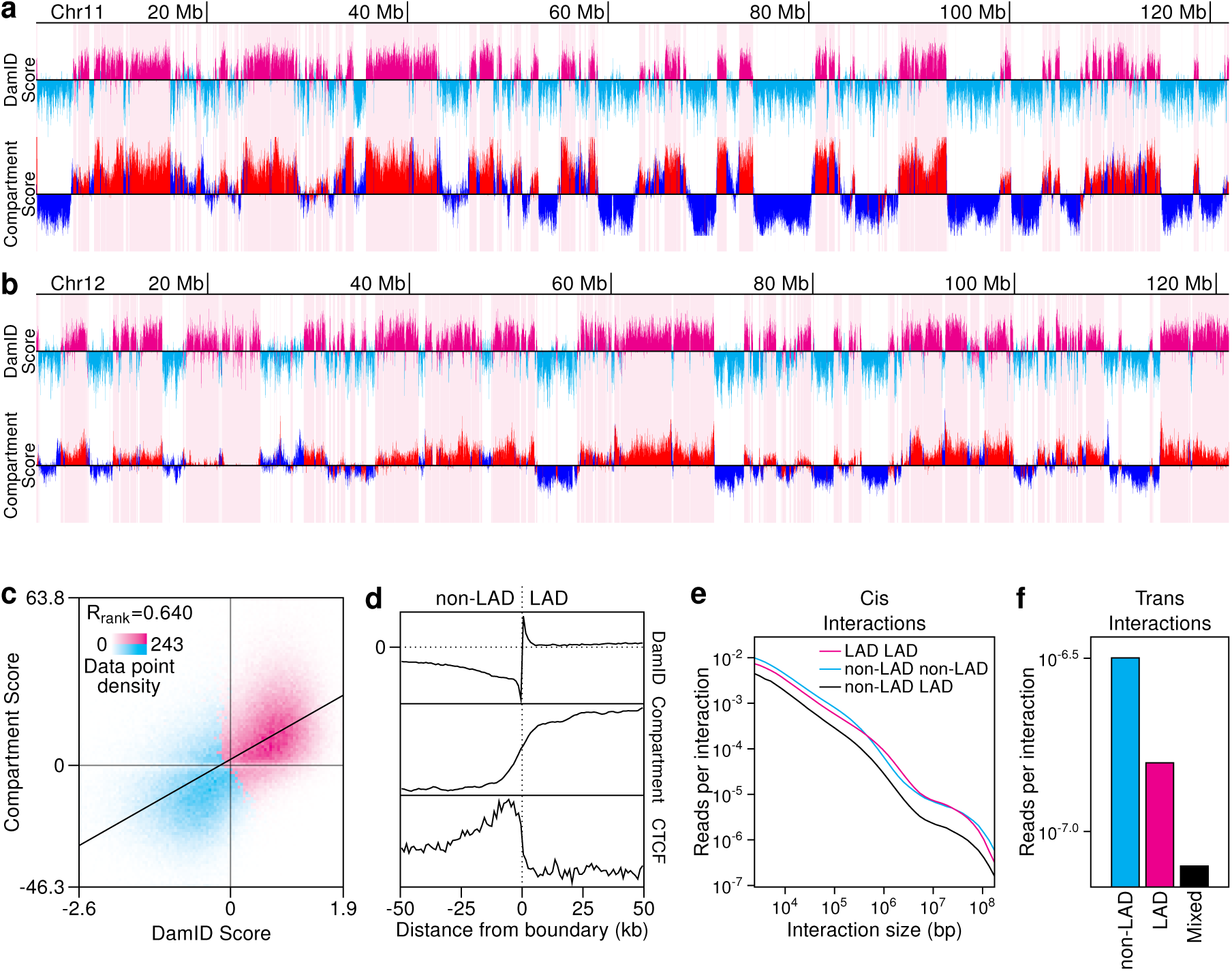
LAD structures captured by both local and chromosome-wide metrics from Hi-C data. **a)** LmnB1 DamID and Hi-C compartment score from chromosome 11 in MEF cells. LAD calls and associated data are highlighted in magenta/red (DamID/compartment scores) while data from non-LAD regions are shown in cyan/blue. **b)** LMNB1 DamID and Hi-C compartment score from chromosome 12 in MEF cells. LAD calls and associated data are highlighted in magenta/red (DamID/compartment scores) while data from non-LAD regions are shown in cyan/blue. **c)** Genome-wide correlation between DamID and compartment scores. Data are partitioned into a 100 by 100 grid with intensity indicating data density and color showing whether the majority of the bins data points are in LADs (magenta) or not (cyan). **d)** Feature profiles anchored at all boundaries of LADs of size 100 kb or greater (excluding chromosome X) and oriented from non-LAD (left) to LAD (right). Profiles consist of data within 100 kb of each boundary binned in 1 kb intervals.**e)** Intra-chromosomal interaction frequencies as a function if interaction size parsed by LAD state. **f)** Inter-chromosomal interaction frequencies parsed by LAD state.

**Supplemental Figure 6:**
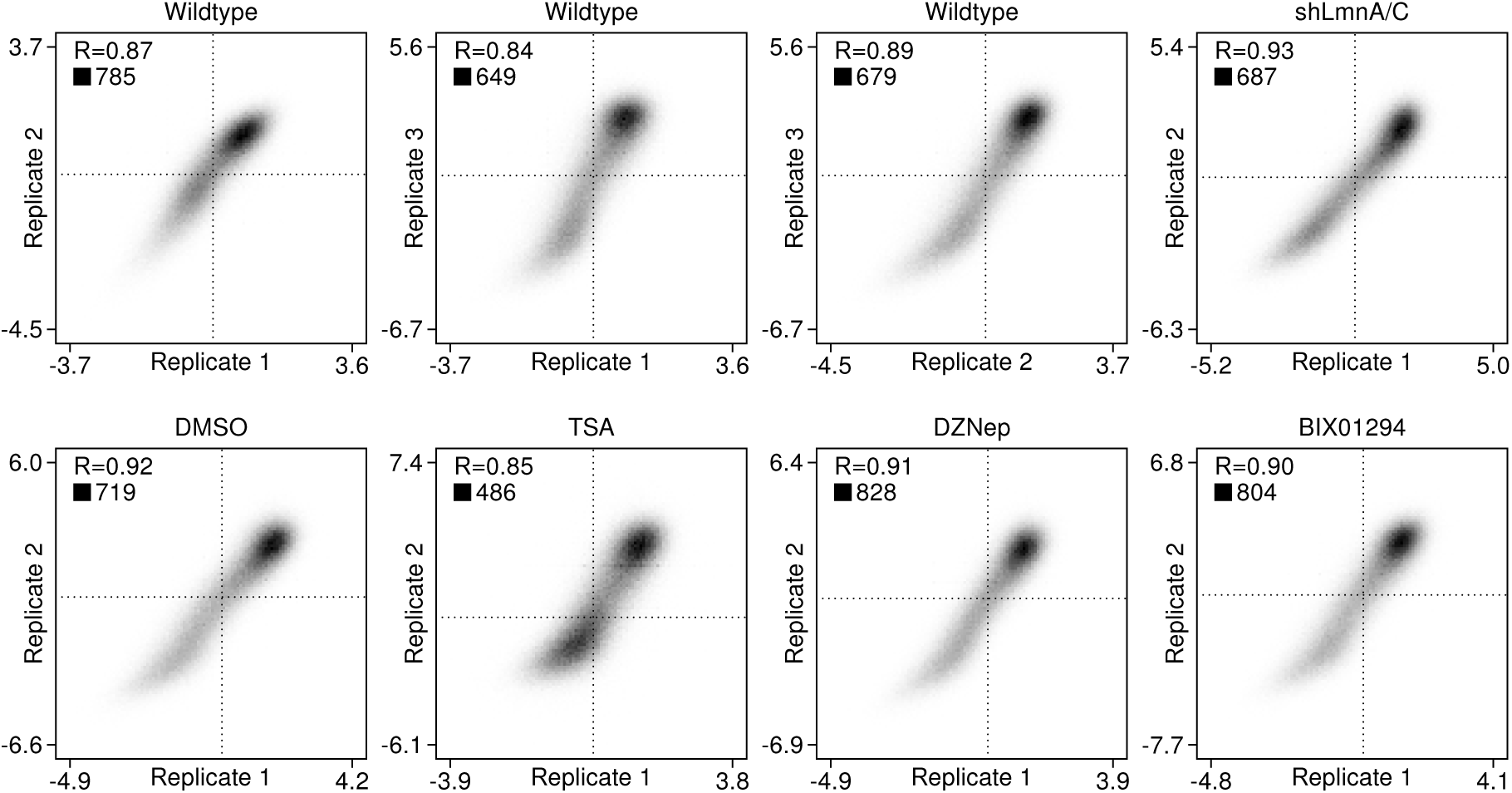
Correlation of DamID runs. Pairwise comparison of replicate LmnB1 DamID scores within each experimental condition. For each comparison, the Pearson correlation coefficient (R) is shown.

**Supplemental Figure 7:**
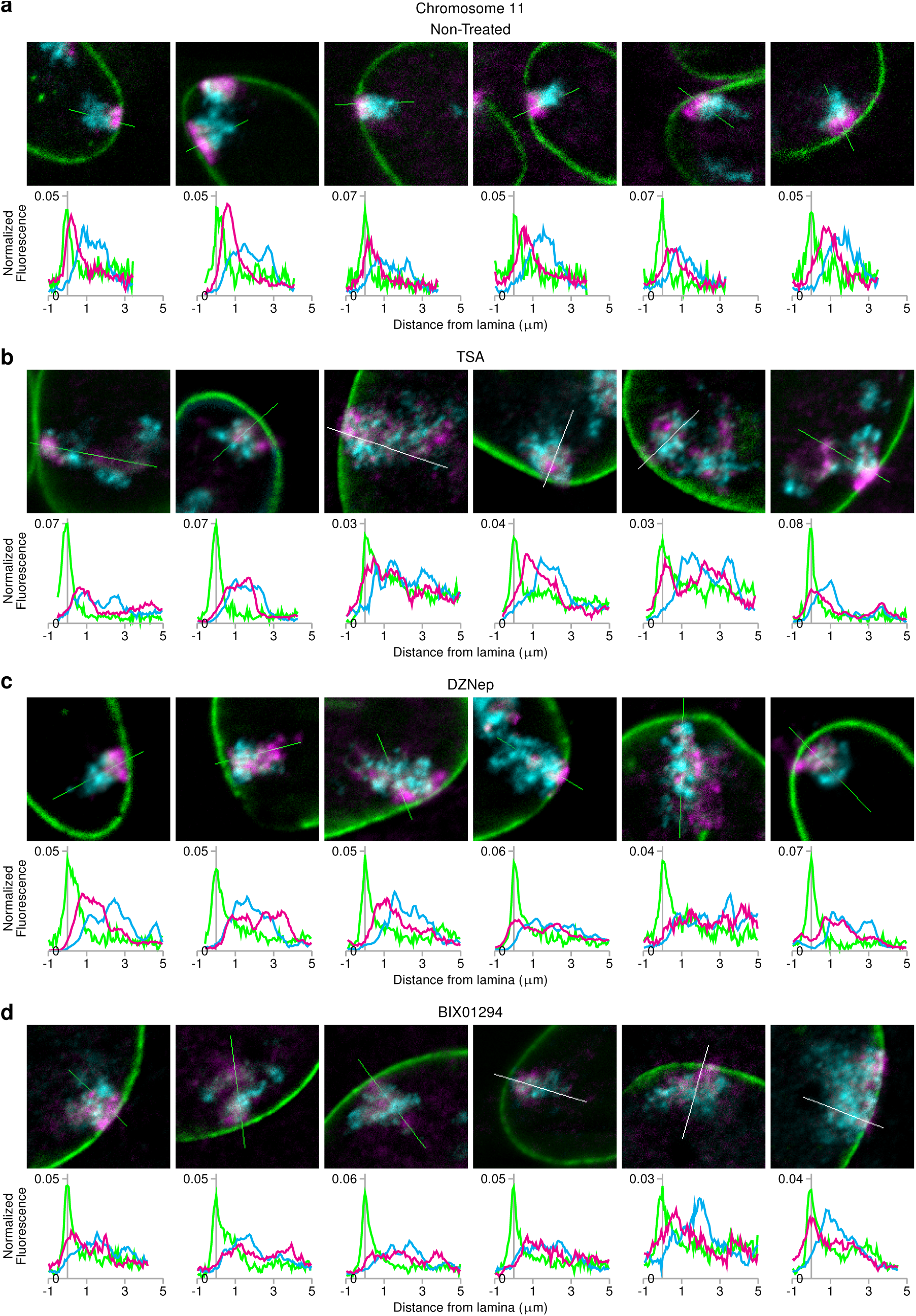
Chromosome 11 conformation profiles after epigenetic perturbation. Example single line scans are presented for **a)** non-treated **b)** TSA-treated **c)** DZNep-treated **d)** and BIX01294-treated. Region measured is indicated over the chromosome territory (overlay line, top) and plotted distribution of LAD, non-LAD, and LmnB1 are shown below (magenta, cyan, and green lines, respectively). All images are shown at the same magnification. All graphs include measurements to 5 µm from the lamina

**Supplemental Figure 8:**
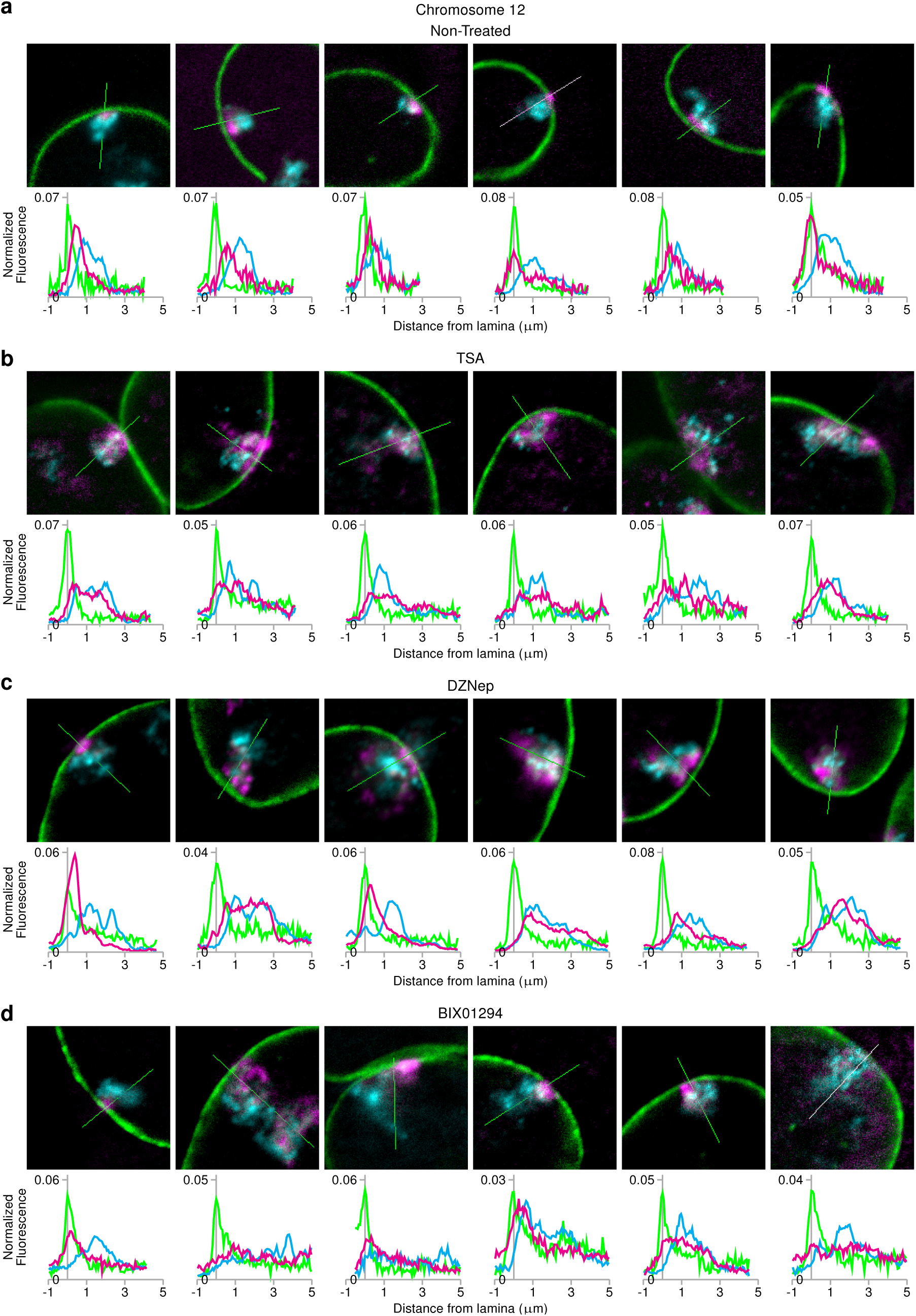
Chromosome 12 conformation profiles after epigenetic perturbation. Example single line scans are presented for **a)** non-treated **b)** TSA-treated **c)** DZNep-treated **d)** and BIX01294-treated. Region measured is indicated over the chromosome territory (overlay line, top) and plotted distribution of LAD, non-LAD, and LmnB1 are shown below (magenta, cyan, and green lines, respectively). All images are shown at the same magnification. All graphs include measurements to 5 µm from the lamina

**Supplemental Figure 9:**
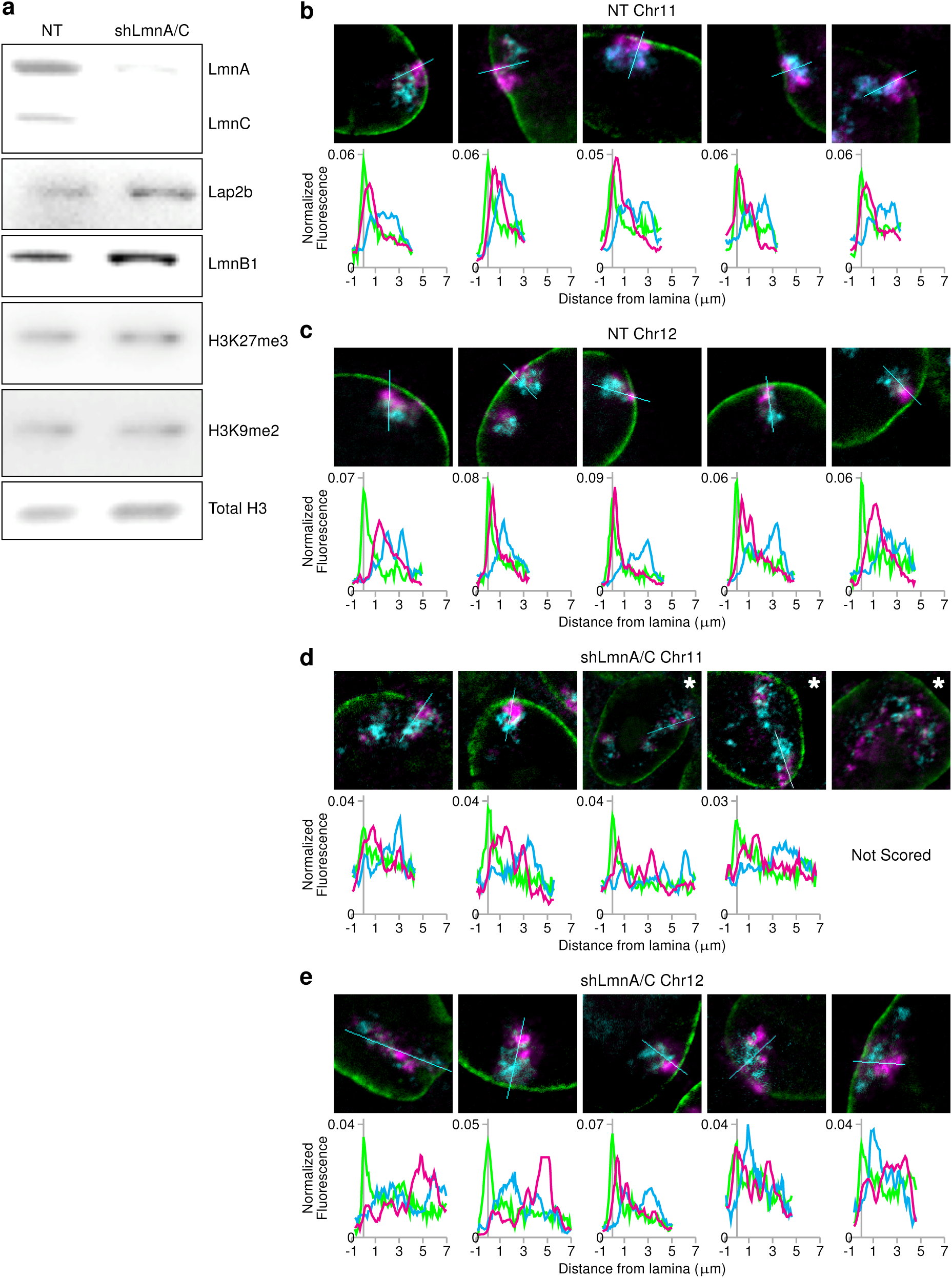
Chromosomes 11 and 12 conformation profiles after shRNA-mediated LmnA/C knockdown. **a)** Immunoblot for the indicated proteins or histone modifications in NT and shLmnA/C cells. Example single line scans are presented for **b)** non-treated (NT) chromosome 11, **c)** NT chromosome 12, **d)** shRNA-mediated LmnA/C knockdown (shLmnA/C) chromosome 11, and **e)** shLmnA/C chromosome 12. Region measured is indicated over the chromosome territory (overlay line, top) and plotted distribution of LAD, non-LAD, and LmnB1 are shown below (magenta, cyan, and green lines, respectively). Images are shown at the same magnification with the exception of starred images which are 2X zoomed out to encompass the measured territory. All graphs include measurements to 7 µm from the lamina.

